# A simple, robust, and low-cost method to produce the PURE cell - free system

**DOI:** 10.1101/420570

**Authors:** Barbora Lavickova, Sebastian J. Maerkl

**Affiliations:** Institute of Bioengineering, School of Engineering École Polytechnique Fédérale de Lausanne Lausanne, Switzerland

**Keywords:** synthetic biology, protein expression, cell-free transcription and translation, PURE system

## Abstract

We demonstrate a simple, robust, and low-cost method for producing the PURE cell-free transcription-translation system. Our OnePot PURE system achieved a protein synthesis yield of 156 *µ*g/mL at a cost of 0.09 USD/*µ*L, leading to a 14-fold improvement in cost normalized protein synthesis yield over existing PURE systems. The OnePot method makes the PURE system easy to generate and allows it to be readily optimized and modified.

## Introduction

Cell-free transcription-translation systems have become popular for molecular engineering (*1*–*6*). Cell-free systems can be categorized into two main classes: cell extract and recombinant systems. Cell extracts are highly functional but complex and undefined cell-free systems. In 2001, Shimizu *et al.* demonstrated that a defined cell-free system called the “PURE” system (protein synthesis using recombinant elements) could be reconstituted from purified recombinant components (*7*). Because of its defined and minimal nature, PURE is an appealing choice for biological systems engineering. The PURE system has been used for genetic network engineering (*2*), recombinant DNA replication (*8*), molecular diagnostics (*9*), therapeutics (*10*), and educational kits (*11*). The PURE system also represents a viable starting point for generation of an artificial cell (*12, 13*) and its composition has been optimized (*14, 15*) and extended (*16*) to achieve higher functionality.

Unfortunately, producing PURE is an arduous and costly process, requiring 36 individual medium to large-scale protein purifications. PURE is now commercially available (PURExpress, New England Biolabs (NEB)), but the high-cost of the commercial system at 1.36 USD/*µ*L still limits its use. Although NEB provides a few different formulations of the PURE system, the commercial system can’t be customized or optimized by the user, and the precise formulation of the commercial PURE system is not publicly available. It was recently demonstrated that the PURE system could be produced using synthetic microbial “consortia” (TraMOS PURE) (*17*), which simplified the process of making PURE by co-expressing multiple protein components in a single *E. coli* clone combined with co-culturing of multiple strains. TraMOS PURE achieved only a ∼20% protein yield compared to the commercial PURExpress and production cost was reduced from 1.36 USD/*µ*L to 0.96 USD/*µ*L. An earlier approach used MAGE to His-tag most PURE protein components in their endogenous locus and co-purified them from 6 strains to generate an ensemble PURE system (ePURE) (*18*). The approach led initially to only minimal protein synthesis activity, and an optimized ePURE system ultimately reached a 11% protein yield compared to the original PURE system (*7*). Shephard *et al.* cloned 30 PURE protein components onto 3 separate plasmids for simplified and low-cost generation of the PURE system (*19*). Upon optimization, this PURE 3.0 system reached protein synthesis levels comparable to the commercial PUREfrex kit (GeneFrontier, Chiba, Japan). As multiple proteins are being expressed and purified from a single *E. coli* clone in all three of these approaches it is not possible to rapidly modify protein levels or omit proteins from the PURE system, which is a critical feature for further PURE system development and optimization.

Here we present a simple, robust, and low-cost method for producing the PURE system. Our method co-cultures (*20*) and induces all 36 protein producing *E. coli* clones in a single flask followed by a single Ni-NTA purification. Our “OnePot” method produces PURE at a cost of 0.09 USD/*µ*L and a protein synthesis capacity of 156 *µ*g/mL, which is as high as the commercial PURE system. OnePot PURE production reduces the cost per microliter to 6% compared to the commercially available PURExpress from NEB (1.36 USD/*µ*L). A single batch prepares enough proteins for a total of 15 mL of PURE which is sufficient material for ∼ 1,500 10 *µ*L reactions and can be generated together with ribosomes in 4 days. The method produces consistent PURE across different batches and allows the rapid optimization of individual PURE protein components.

## Results and Discussion

The PURE system consists of several different components (*7*), that can be separated into three main categories: proteins (transcription, translation, and energy regeneration), ribosomes, and small molecule components (salts, buffers, NTPs, creatine phospate, and folinic acid). In this work, we developed a “OnePot” method for the preparation of all 36 protein components using a single mixed co-culture and Ni-NTA affinity purification step to simplify the process and decrease the cost of the PURE system. All 36 *E. coli* expression clones are cultured individually in small volumes overnight, which are then combined to inoculate a single 500 mL culture. The mixed culture is allowed to outgrow and is induced, followed by pelleting, lysis and loading of the lysate onto a Ni-NTA column for protein purification. To keep the final cost of the PURE system as low as possible, we also prepared ribosome and energy solutions (Fig. 1a, Supplementary Fig. S1, S2). The entire process of OnePot PURE system preparation, including protein and ribosome purification and energy solution preparation, requires 4 days with 20 hours of hands-on time (Supplementary Table S1, S2, S3). To date no method has been presented in which all non-ribosomal PURE proteins were prepared using a single co-culture and purification step (*17, 18*). Moreover, other simplified protocols resulted in low protein synthesis activity as compared to the original PURE system (*17*).

**Figure 1:**
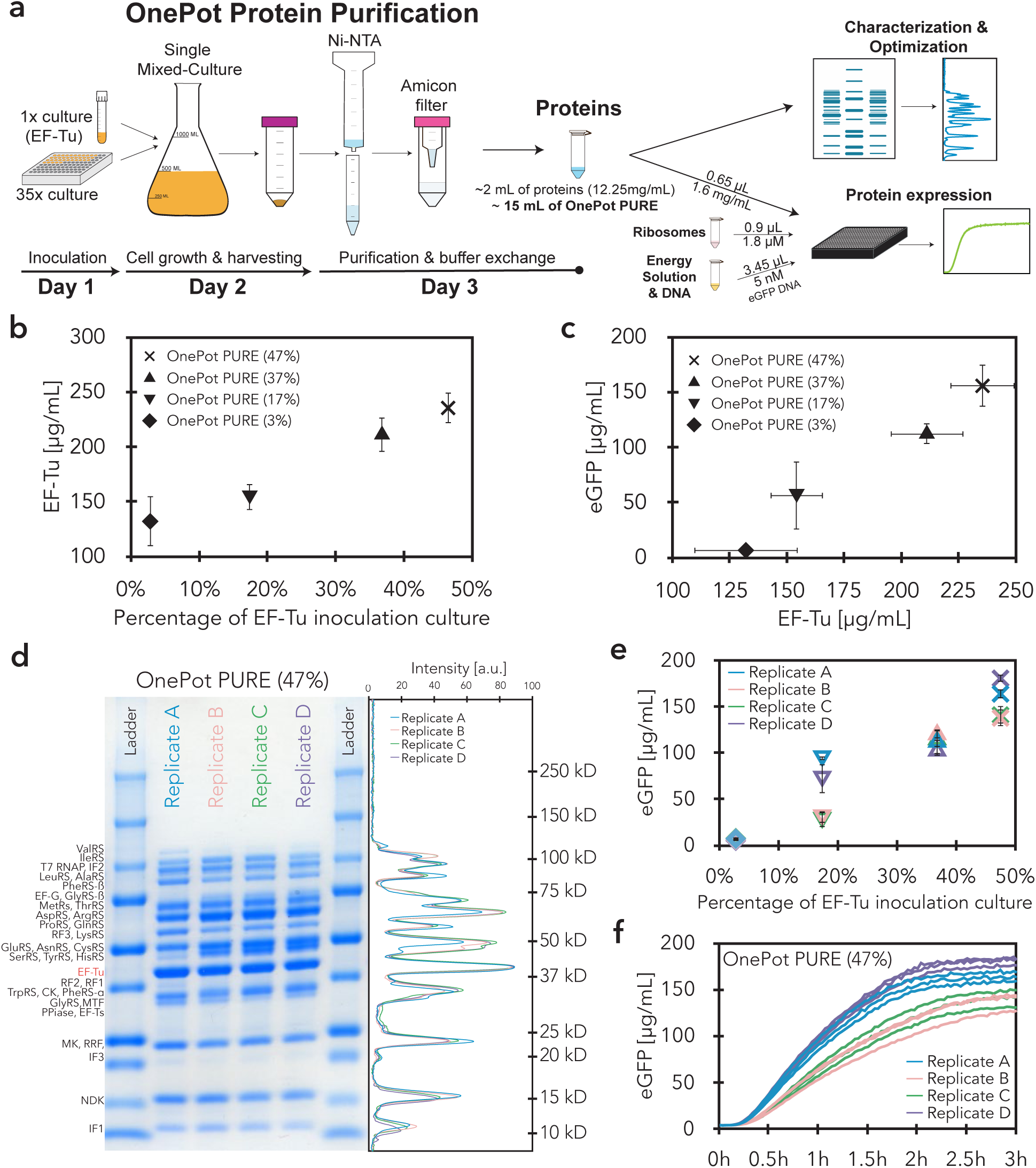
OnePot PURE preparation and optimization. **(a)** All 36 PURE protein components were produced using the OnePot method, which consists of a single co-culture and a single Ni-NTA affinity purification. Different OnePot systems were produced by varying the ratio of inoculation culture EF-Tu with respect to the 35 remaining inoculation cultures, and characterized using SDS-PAGE gels and eGFP expression. **(b)** Concentration of EF-Tu in OnePot PURE reactions derived from SDS-PAGE gel analysis, as a function of relative volume ratios of the EF-Tu inoculation culture in a co-culture. Each data point represents four biological replicates (mean ± s.d.). **(c)** *In vitro* eGFP expression activity after 3h plotted against concentration of EF-Tu in OnePot PURE reactions. Measurements on the x-axis represent biological replicates, and y-axis measurements represent four biological replicates with three technical replicates. Error bars represent s.d. **(d)** Coomassie blue-stained SDS-PAGE gel of four OnePot PURE (EF-Tu 47%) replicates. In the right panel, intensities of the different replicates are plotted with molecular weight standards (kDa). **(e)** *In vitro* eGFP expression activity after 3 h plotted against relative inoculation volume ratios of EF-Tu. Each data point represents a single biological replicate with three technical replicates; error bars represent s.d. of the technical replicates. **(f)** Time course of *in vitro* eGFP expression with OnePot PURE (EF-Tu 47%). Each line represents a technical replicate.

We explored whether it is possible to adjust the protein component ratios in the OnePot PURE system simply by varying the ratios of the inoculation culture volumes added to the mixed co-culture (Supplementary Table S4). Besides ribosomal proteins, elongation factor thermo unstable (EF-Tu) is one of the most abundant proteins in rapidly growing *E. coli* (*21*) and it was shown to be one of the key factors for *in vitro* protein synthesis (*14*). Previous work showed that PURE is otherwise relatively robust to changes in protein concentrations as demonstrated by experimental work where PURE protein components were titrated (*14, 22*) and computational modeling (*23*). Additionally, over 50% of the HomeMade PURE protein components consist of EF-Tu (Supplementary Table S5). Hence, we decided to optimize our OnePot PURE system with a particular focus on this translation factor.

We varied the relative volume of the EF-Tu inoculating culture with respect to the 35 remaining inoculation cultures to generate ratios of 3%, 17%, 38%, and 47%. The 3% ratio corresponds to 100 *µ*L of all 36 inoculation cultures, including EF-Tu, being added to the mixed co-culture (Supplementary Table S4). As can be seen from gels and corresponding analysis, larger percentages of the EF-Tu strain in the co-culture led to higher absolute levels of EF-Tu in the OnePot protein system (Fig. 1b, Supplementary Fig. S3, S4). Increased concentrations of EF-Tu also gave rise to higher protein expression yields (Fig. 1c). We could therefore show that it is possible to modify the ratio of an individual PURE protein component simply by varying the initial inoculation ratio of the corresponding strain, and that the OnePot PURE system gave rise to high protein expression yields.

It has been thought that precise control over the PURE system composition is required to achieve reproducible, and high protein expression yields and it has been suggested that a simple one-pot method would not be a viable option for robustly generating the PURE system (*17*). However, we observed that variations in overnight culture densities (Supplementary Fig. S5) did not lead to substantial differences in OnePot PURE protein content (Fig. 1d, Supplementary Fig. S3, S6c-e). We observed high protein expression robustness across four biological replicates, especially for the 38% and 47% EF-Tu formulations, with coefficients of variation (CV) of 8% and 12%, respectively (Fig. 1e, f). In comparison, the CV for a technical replicate of PURExpress and HomeMade PURE were 5% and 12%, respectively. To avoid significant total protein concentration differences across replicates, we adjusted the concentration of the protein mixture to 1.6 mg/mL in the final reaction. This optimal concentration was chosen based on titrations of OnePot PURE (47% EF-Tu) replicate A (Supplementary Fig. S7).

We compared the protein composition of our OnePot PURE system to the commercially available PURExpress (NEB) and our HomeMade PURE system prepared based on the Shimizu protocol with minor adjustments (*7*). From gels and mass spectrometry (MS) we determined that the overall composition of the PURExpress and HomeMade PURE systems were quite similar to one another as expected (Fig. 2a, Supplementary Fig. S6d). Both PUR-Express and HomeMade PURE had a higher relative percentage of EF-Tu and a lower total protein concentration (1 mg/mL for HomeMade PURE) than OnePot PURE. The relative intensities of individual proteins in the OnePot PURE deviated from the PURExpress and HomeMade PURE standards although the protein expression yield of the OnePot PURE system (47% EF-Tu) was similar to PURExpress, 1.6 times higher than our HomeMade PURE and 5 times higher than TraMOS (Fig. 2b).

**Figure 2:**
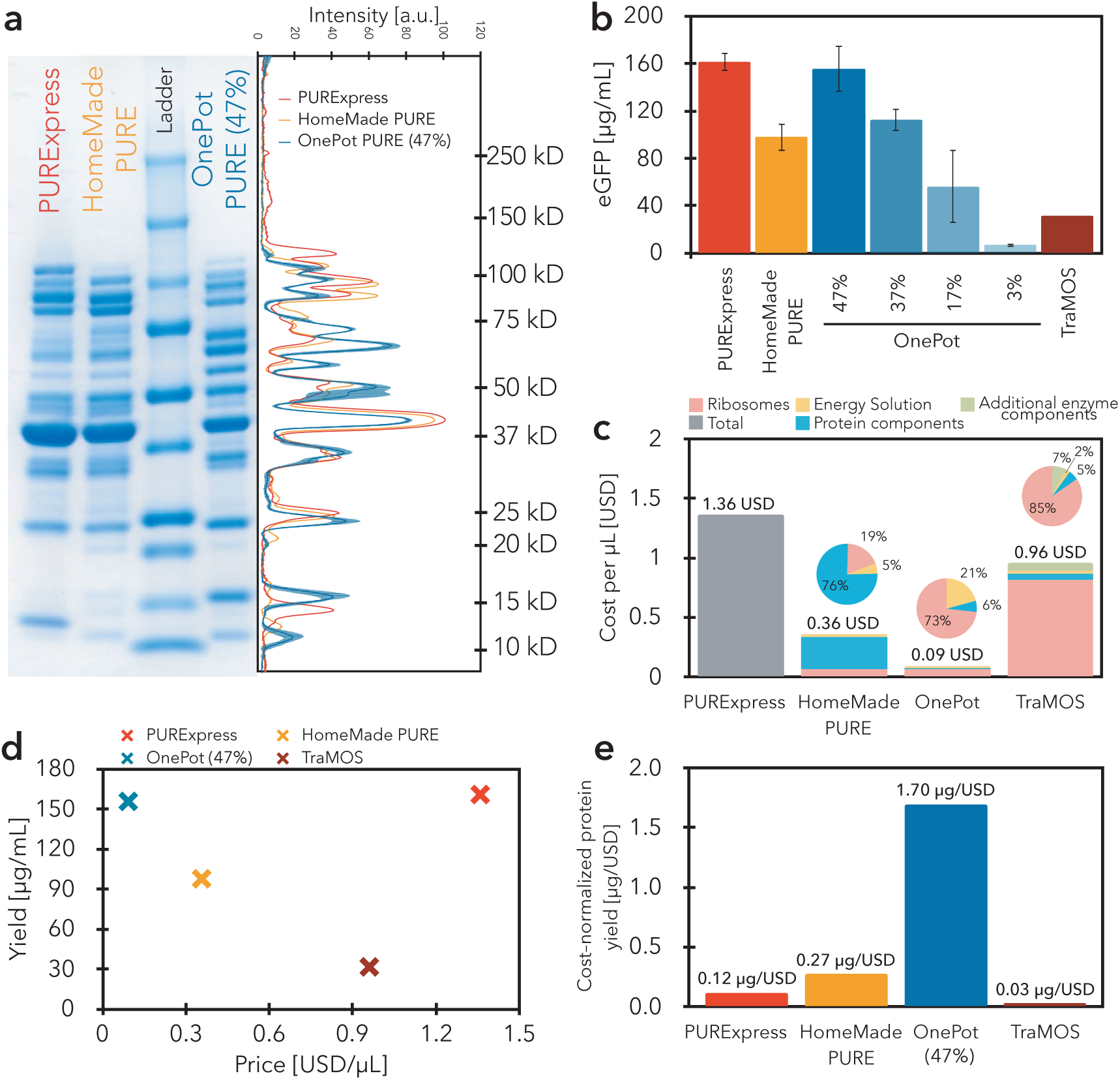
OnePot PURE comparison to existing PURE systems. **(a)** SDS-PAGE gel of PURExpress, HomeMadePURE, OnePot PURE (EF-Tu 47%, replicate A). In the right panel, intensities of different replicates are plotted with molecular weight standards (kDa). **(b)** Comparison of eGFP expression activity (after 3 h) of different PURE systems. The different systems were tested in the same conditions except for TraMOS where the reported value was used (*17*). **(c)** Price comparison of the different PURE systems. Calculations are detailed in Supplementary Tables S1, S2, S3, Supplementary Table S6, S7. **(d)** Yield of the different PURE systems plotted against their price per *µ*L. Mean values of the eGFP expression yield were plotted. **(e)** Cost-normalized yield of the different PURE systems. The mean value of the eGFP expression yield was used for the calculations.

Moreover, we compared expression levels of different proteins in PURExpress and OnePot PURE (47% EF-Tu). Based on SDS-PAGE gels of proteins labeled with FluoroTect GreenLys tRNA, we reached similar levels of expression in PURExpress and OnePot PURE for eGFP (26.9 kDa), T3 RNAP (98.8 kDa), *β*-galactosidase (116.5 kDa) and trehalase (63.7 kDa) (Supplementary Fig. S8). We were not able to separate bands for DHFR (18 kDa) as it co-migrated with FluoroTect GreenLys tRNA bands. However, we were able to distinguish the expected bands for all four proteins on a Coomassie-stained gel (Supplementary Fig.S8b). Activity assays for *β*-galactosidase and trehalase (Supplementary Fig. S9) showed that the synthesized proteins were functional. We also synthesized a zinc-finger transcription factor demonstrating functional repression of deGFP (*24, 25*), and achieved comparable fold-repression levels in PURExpress and OnePot PURE supplemented with *E. coli* RNA polymerase and *σ*70 factor (Supplementary Fig. S9).

Since PURE systems are prepared by affinity chromatography, a certain amount of contaminants can be expected in the systems. To approximate the amount of protein contamination present we analyzed PURExpress, HomeMade PURE and OnePot PURE by LC-MS/MS. The percentage of contaminants was estimated based on total independent spectral counts, which correlated with the amount of protein present in the sample (Supplementary Fig. S6a, b). Our OnePot method gave rise to a similar amount of contamination as in-house prepared HomeMade PURE (Supplementary Fig. S6e). The amount of contamination across all OnePot PURE (EF-Tu 47%) replicates was 12.6 ± 1.5% and for the HomeMade PURE 11.3%. PURExpress had a lower level of contamination of 4.5%. Moreover, the contaminants present across the different PURE systems are similar; more than 50% of contaminants present in OnePot PURE are present in HomeMade PURE as well (Supplementary Excel file for LC-MS/MS). Moreover, many of these contaminants are well-known His-tag based purification contaminants (*26*). The main difference is the presence of ribosomal proteins (Supplementary Fig. S6f, Supplementary Excel file for LC-MS/MS), these represent around 40% of the contaminating proteins present in OnePot PURE only. This observation is in agreement with results obtained for TraMOS (*17*). Based on these results the OnePot PURE system achieved similar levels of purity as PURE produced in-house using the standard method.

One of the main factors limiting the use of the PURE system is its high cost. We performed a detailed cost analysis of different PURE systems: two systems prepared from individually purified protein components (PURExpress and HomeMade PURE), as well as two systems prepared from batch cultures and pooled purifications (OnePot and TraMOS) (Fig. 2c, Supplementary Table S1, S2, S3, Supplementary Table S6, S7). The commercial PURExpress is the most expensive at a cost of 1.36 USD/*µ*L followed by TraMOS (0.96 USD/*µ*L), HomeMade PURE (0.36 USD/*µ*L), and OnePot PURE (0.09 USD/*µ*L). For the HomeMade PURE and TraMOS preparations, cost originates primarily from protein components and ribosomes. The OnePot approach reduces the cost of the non-ribosomal protein components to negligible levels and relies on ribosome purification to further reduce cost. Combining in-house ribosome purification with bulk purification of non-ribosomal proteins is thus a general strategy to reduce cost. In-house ribosome purification does not only reduce the price by almost 16-fold as compared to using commercial ribosomes, but also allows for higher ribosome concentrations in the PURE system. The standard ribosome purification protocol used in this work is simple and robust. We compared a total of six ribosome preparations purified over a period of 11 months (Supplementary Fig. S10a) showing similar expression levels in OnePot PURE for all batches, demonstrating the robustness of the purification process as well as long-term stability of the purified ribosomes (Supplementary Fig. S10b). Moreover, in case ultracentrifugation is not accessible, His-tag purification of ribosomes could be a viable alternative (*18, 27*). OnePot PURE substantially outperformed all other systems when directly comparing protein synthesis yield and cost per microliter (Fig. 2d), achieving a cost normalized protein yield of 1.70 *µ*g/USD compared to 0.27 *µ*g/USD for HomeMade PURE, 0.12 *µ*g/USD for PURExpress, and 0.03 *µ*g/USD for TraMOS (Fig. 2e).

We demonstrated that it is possible to robustly produce a highly functional PURE system at low cost using a practical single batch culture and purification approach. Previous approaches such as ePURE, TraMOS, and PURE 3.0 all expressed multiple proteins within a single host to simplify downstream purification. TraMOS further combined this concept with co-culturing of multiple strains but failed to produced highly functional PURE from a 34 strain co-culture in which each strain expressed a single protein. Here we show that 36 strains can be successfully co-cultured, eliminating the need to co-express multiple proteins in a single host. This in turn makes it possible to rapidly adjust the formulation of the resulting PURE mix which would require tedious and time-consuming cloning steps with the previous methods. The OnePot PURE system described here achieved a protein synthesis yield of 156 *µ*g/mL at a cost of 0.09 USD/*µ*L. At 1.7 *µ*g/USD the cost normalized protein synthesis yield is over a magnitude higher than the commercial PURE system and substantially higher than TraMOS. We also showed that it is possible to adjust and optimize the OnePot PURE system by varying the inoculation fraction of an individual strain. This simple, low-cost, and robust protocol for producing the PURE system should broaden access to the technology and enable new applications which hitherto were not feasible due to the high cost and complexity of producing the PURE system.

## Methods

### *Escherichia coli* strains, plasmids, and linear DNA templates

*E. coli* BL21(DE3) and M15 strains were used for protein expression. All plasmids encoding PURE proteins used in this work were originally obtained from Y. Shimizu (RIKEN Quantitative Biology Center, Japan). Genes coding for MK and PPiase were originally cloned in pET29b vectors with kanamycin resistance. To establish a OnePot system, we used CPEC assembly (Circular Polymerase Extension Cloning) (*28*) to clone a DNA fragment amplified from pET29b vectors containing MK and PPiase genes as well as the T7 promoter, RBS, and T7 terminator, into a pET21a vector containing ampicillin resistance. The primer sequences used are listed in Supplementary Table S8. A list of the PURE proteins with their corresponding gene, vector and reference number are given in Supplementary Table S9. *E. coli* A19 (Coli Genetic Stock Center, CGSC#: 5997) was used for ribosome purification.

Linear template DNA for *in vitro* eGFP synthesis was initially prepared by extension PCR from a pKT127 plasmid as described (*2*) and cloned into a pSBlue-1 plasmid. The DNA fragment used for PURE system characterization was amplified from this plasmid by PCR. DNA templates coding for trehalase, *β*-galactosidase and T3 RNA polymerase were amplified from *E. coli* MG1655Z1 genome, ZIKV Sensor 27B LacZ (Addgene plasmid # 75006) (*9*) and BBa K346000 (Registry of Standard Biological Parts), respectively, by extension PCR. Primer sequences are listed in Supplementary Table S10. For DHFR expression the control template supplied with PURExpress was used. DNA templates for Zinc-fingers (Supplementary Table S11) were prepared as described (*25*). DNA fragments were purified using DNA Clean and Concentrator-25 (Zymo Research). DNA was eluted in nuclease-free water instead of elution buffer.

### Buffers used for protein and ribosome purification

All buffers used in this work are listed in Supplementary Table S12. All buffers were filtered (Flow Bottle Top Filters, 0.45 *µ*m aPES membrane) and stored at 4°C. 2-mercaptoethanol was added immediately before use.

### OnePot protein preparation

Lysogeny broth (LB) used for OnePot protein component preparation was supplemented with 100 *µ*g/mL ampicillin and all cultures were grown at 37°C, 260 RPM. To allow for fast and easy inoculation, the different strains were stored as a glycerol stock in a single 96 well microplate. All overnight cultures were inoculated by a 96-well replicator (VP 408FS2AS, V & P Scientific), except for the EF-Tu strain, and grown in 0.3 mL of LB in a deep-well microplate (96 wells, void volume 1.5 mL). The strain expressing EF-Tu was grown in 3 mL of LB in a standard 14 mL culture tube. Overnight cultures (in total 3.6 mL) were used to inoculate 500 mL of LB media in a 1 L baffled flask. The exact composition of the inoculation cultures for different OnePot systems are given in Supplementary Table S4. Cells were grown 2 h before induction with 0.1 mM IPTG for 3 h, then harvested by centrifugation (4,000 RPM, 10 min, 4°C) and stored at −80°C overnight. Cells were resuspended in 7.5 mL buffer A and lysed by sonication on ice (Vibra cell 75186 and probe tip diameter: 6 mm, 4 × 20s:20s pulse, 70% amplitude). Cell debris was removed by centrifugation (15,000 RPM, 20 min, 4°C). The supernatant was mixed with 2 mL of equilibrated resin, prepared as described below, and incubated for 3 h, at 4°C. After the incubation, unbound lysate was allowed to flow through the column. The column was washed with 25 mL of a wash buffer (95% buffer A, 5% buffer B) and eluted with 5 mL of elution buffer (10% buffer A, 90% buffer B). Instead of dialysis, buffer exchange was done using a 15 mL Amicon Ultra filter unit with a 3 kDa molecular weight cutoff (Merck). All centrifugation steps were performed at 4,000 RPM and 4°C. The elution fraction was diluted with 25 mL of HT buffer and concentrated to 1 mL (2 × 60 min). The concentrated sample was then diluted with 10 mL of HT buffer, concentrated to 1.5 mL (60 min), and mixed with 1.5 mL of stock buffer B. The protein solution was then concentrated (14,000 RPM, 30 min, 4°C) using a 0.5 mL Amicon Ultra filter unit with a 3 kDa molecular weight cutoff (Merck) and stored at −80°C. Total protein concentration in the OnePot protein mixture was determined using a microplate Bradford protein assay with bovine gamma-globulin as a standard (Bio-Rad). Samples were diluted 1:25 and 5 *µ*L of the diluted sample was mixed with 250 *µ*L of Bradford reagent. Absorbance at 595 nm was measured using a SynergyMX platereader (BioTek). The OnePot protein mixture was then adjusted to a concentration of 12.25 mg/mL.

### HomeMade PURE protein preparation

Proteins were prepared by Ni-NTA gravity-flow chromatography. The LB medium used was supplemented with 100 *µ*g/mL of ampicillin and/or 50 *µ*g/mL of kanamycin (Supplementary Table S5), and all cultures were grown at 37°C, 250 RPM. Overnight cultures were grown in 3 mL of LB. Each strain was then individually inoculated in a flask with 2 L of LB. Cells were grown 2 h before induction with 0.1 mM of IPTG for 3 h, then harvested by centrifugation and stored at −80°C overnight. The cells were resuspended in 30 mL of buffer A and lysed by sonication on ice (Vibra cell 75186 and probe tip diameter: 6 mm, 8 × 20s:20s pulse, 70% amplitude). Cell debris was removed by centrifugation (25,000 RCF, 20 min, 4°C). The supernatant was mixed with 2-3 mL of equilibrated resin (described below), and incubated for 1-2 h, at 4°C. After the incubation, unbound lysate was allowed to flow through the column. The column was washed with 30 mL of a wash buffer (95% buffer A, 5% buffer B) and eluted with 15 mL of an elution buffer (10% buffer A, 90% buffer B). The elution fraction was dialysed against HT buffer (2×) and stock buffer and stored at −80°C. Protein concentrations were estimated by absorbance at 280 nm and calculated protein extinction coefficients. When a higher protein concentration was required, the protein solution was concentrated using a 0.5 mL Amicon Ultra filter unit (Merck).

### Ni-NTA resin preparation and regeneration

2 mL IMAC Sepharose 6 FF (GE Healthcare) was pipetted into Econo-Pac chromatography columns (Bio-Rad), and charged with 15 mL of 100 mM nickel sulfate solution. The charged column was washed with 50 mL of DEMI water and equilibrated with 35 mL of buffer A. After protein purification, columns were regenerated with 10 mL of buffer containing 0.2 M EDTA and 0.5 M NaCl, and washed with 30 mL of 0.5 M NaCl, followed by 30 mL of demineralized water, and stored in 20% ethanol at 4°C.

### OD600 measurement

OD600 measurements of over-night cultures were measured on a 96-well plate with tenfold dilutions (20 *µ*L of over-night culture in 180 *µ*L of LB) using a SynergyMX platereader (BioTek). The background (OD600 of 200 *µ*L of LB) was subtracted from all samples.

### Ribosome purification

Ribosomes were prepared from *E. coli* A19 by hydrophobic interaction chromatography (HIC) and sucrose cushion buffer ultracentrifugation as described previously with slight modifications (*29, 30*). *E. coli* A19 strain was grown overnight in 100 mL of LB media at 37°C. 2 × 30 mL of the overnight cultures was used to inoculate 2 × 2 L of LB. Cells were grown at 37°C, 250 RPM to exponential phase (3-4 h, OD600 = 0.6-0.8), harvested by centrifugation (4,000 RCF, 20 min, at 4°C), resuspended in 50 mL suspension buffer and stored at −80°C. The resuspended cells were lysed by sonication on ice (Vibra cell 75186 and probe tip diameter: 6 mm, 12 × 20s:20s pulse, 70% amplitude). The cell debris was removed by centrifugation (20,000 RCF, 20 min, at 4°C). The recovered fraction was mixed with the same amount of high salt suspension buffer. The precipitate was removed by centrifugation (20,000 RCF, 20 min, at 4°C) and the supernatant was filtrated with a GD/X syringe filter membrane (0.45 mm, PVDF, Whatman).

Ribosomes were purified using a 15 mL (3 × 5 mL HiTrap Butyl HP column (GE Health-care) on Akta Purifier FPLC (GE Healthcare) at 4°C. After the column was equilibrated with 60 mL of buffer C, the prepared lysate solution was loaded onto the column and washed with 45 mL of wash buffer 1 (100% buffer C) followed by 75 mL of wash buffer 2 (80% buffer C, 20% buffer D). Ribosomes were eluted with 60 mL of ribosome elution buffer (50% buffer C, 50% buffer D) followed by 60 mL of final elution buffer (100% buffer D) at a flow rate of 4 mL per minute. All fractions containing ribosomes (absorbance peak at 280 nm during elution with ribosome elution buffer) were pooled together (around 55 mL). The column was recovered by washing with NaOH (1 M) and acetic acid (0.1 M), and stored in 20% ethanol. 14 mL of recovered fraction was overlaid onto 15 mL of cushion buffer in four polycarbonate tubes (void volume: 32 mL). The ribosomes were pelleted by ultracentrifugation (Beckman type SW 32 Ti rotor, 100,000 RCF, 16 h, 4°C). Each transparent ribosome pellet was washed two times with 0.5 mL ribosome buffer and resuspended with a magnetic stirrer in 100 *µ*M of ribosome buffer. To ensure that all the ribosomes are recovered every tube was washed with 100 *µ*M ribosome buffer. The recovered solution was concentrated using a 0.5 mL Amicon Ultra filter unit with a 3 kDa molecular weight cutoff (Merck) by centrifugation (14,000 RCF, 10 min, at 4°C). Ribosome concentrations were determined by measuring absorbance at 260 nm of a 1:100 dilution. An absorbance of 10 for the diluted solution corresponded to a 23 *µ*M concentration of undiluted ribosome solution. Final ribosome solution used for *in vitro* protein synthesis was prepared by diluting the sample to 10 *µ*M. The usual yield is above 0.75 mL of 10 *µ*M ribosome solution.

### SDS-PAGE gels

Proteins were separated by SDS-PAGE using 15-well 4-20% Mini-PROTEAN TGX Precast Protein Gels (Bio-Rad). Gels were stained using Bio-Safe Coomassie stain (Bio-Rad), scanned using an EPSON Perfection V10 scanner and analyzed with ImageJ. In case of all gels containing PURE proteins a mixture 0.625 *µ*L of the adjusted solution was loaded. Concentration of EF-Tu in different PURE systems was determined based on SDS-PAGE gels of EF-Tu with known concentrations (Supplementary Fig. S11). PURE reactions (5 *µ*L) labeled with FluoroTect GreenLys (Promega) were analyzed by SDS-PAGE using 15-well 4-20% Mini-PROTEAN TGX Precast Protein Gels (Bio-Rad) scanned (AlexaFluor 488 settings, excitation: Spectra blue 470nm, emission: F-535 Y2 filter) at Fusion FX7 (Vilber).

### Mass spectrometry

Prior the MS analysis, 15 *µ*L of PURE proteins was subjected to buffer exchange. The samples were diluted to 500 *µ*L in 100 mM ammonium bicarbonate buffer and concentrated by 0.5 mL Amicon Ultra 3 kDa filter unit by centrifugation (14,000 RCF, at 4°C) to 100 *µ*L. This process was repeated three times, with 100 *µ*L of the sample prepared for tryptic digestion and LC-MS/MS analysis. Samples were submitted to tryptic digestion as follows. First, 90 *µ*L of each sample were denaturated by heating for 10 min at 95°C. Then, disulfide bridges were reduced by incubation with tris(2-carboxyethyl)phosphine at 15 mM final concentration for 1 h at 30°C. Cysteine residues were subsequently alkylated for 30 min with iodoacetamide at 20 mM final concentration at room temperature in the dark. Afterwards trypsin was added to the reaction mixture in the ratio 1:50 for overnight digestion. Reaction was quenched by addition of trifluoroacetic acid to 1% final concentration.

Digested samples containing proteolytic peptides were analyzed by LC-MS/MS. 5 *µ*L of each sample were loaded onto a Zorbax Eclipse Plus C18 (1.8 *µ*m, 2.1 x 150 mm) analytical column from Agilent Technology for separation using analytical Dionex Ultimate 3000 RSLC system from Thermo Scientific. The separation was performed with a flow rate of 250 *µ*L/min by applying an effective gradient of solvent B from 5 to 35% in 60 min, followed by column washing and re-equilibration steps. Solvent A was composed of water with 0.1% formic acid, while solvent B consisted of acetonitrile with 0.1% formic acid. The outlet of the chromatographic column was coupled online with the conventional HESI source from Thermo Scientific and eluting peptides were analyzed by high resolution QExactive HF-HT-Orbitrap-FT-MS benchtop mass spectrometer from Thermo Scientific. Analysis was performed in data-dependent manner with 60000 resolution and AGC (automatic gain control) of 3e6 for MS1 scan. MS2 scans were realized in Top10 mode with dynamic exclusion of 30 sec, 15000 resolution, 2 uscans, AGC of 1e5, precursor isolation window of 2 m/z and NCE (normalized collision energy) of 27% for HCD (higher energy collisional dissociation) fragmentation.

Obtained shotgun bottom-up proteomic data were processed with open source TransProteomic Pipeline software (Institute for System Biology, Seattle Proteome Center) using Xtandem! search engine. Peptides were searched against custom database containing all *E.coli* proteins from SwissProt database (Uniprot) together with creatine kinase and adenylate kinase from *Gallus gallus*, inorganic pyrophosphatase from *S. cerevisiae*, T7 RNA polymerase from enterobacteria phage T7 and cationic trypsin from *Bos taurus*. The precursor ion mass tolerance was set to 10 ppm with product ion tolerance of 0.02 Da. Cysteine carbamidomethylation was set as fixed modification, while methionine oxidation, asparagine/glutamine deamidation and N-terminal acetylation were specified as dynamic modifications. The cleavage specificity was set to trypsin with two allowed missed cleavages. 1% false discovery rate (FDR) was allowed with minimum peptide length of 7 amino acids and minimum 2 peptides per protein.

### Energy solution preparation

Energy solution was prepared as described previously with slight modifications (*30*). 2.5× energy solution contained 0.75 mM of each amino acid, 29.5 mM of magnesium acetate, 250 mM of potassium glutamate, 5 mM of ATP and GTP, 2.5 mM CTP, UTP, and DTT (Dithiothreitol), 130 U_*A*260_/mL of tRNA, 50 mM of creatine phospate, 0.05 mM of folinic acid, 5 mM of spermidine, and 125 mM of HEPES.

### *In vitro* protein expression and functional assays

HomeMade or OnePot PURE reactions (5 *µ*L) were established by mixing 2 *µ*L of 2.5x energy solution, 0.9 *µ*L of 10 *µ*M ribosomes (final concentration: 1.8 *µ*M), 0.65 *µ*L of PURE proteins (HomeMade or OnePot solution), DNA template and brought to a final volume of 5 *µ*L with addition of water. PURExpress reactions (5 *µ*L) were established by mixing 2 *µ*L of solution A, 1.5 *µ*L of solution B, DNA template and brought to 5 *µ*L with water.

All reactions measuring eGFP expression levels were prepared as described above with eGFP linear template at a final concentration of 5 nM and incubated at 37°C at constant shaking for 3 h, and measured (excitation: 488 nm, emission: 507 nm) on a SynergyMX platereader (BioTek). Absolute eGFP concentrations were determined from a standard curve (Supplementary Fig. S12).

Reactions expressing other proteins were prepared as described above and supplemented with 0.2 *µ*L FluoroTect GreenLys (Promega). DNA templates at a final concentration of 5 nM, except DHFR which was supplied at a concentration of 10 ng/*µ*L, were used. The reactions were incubated at 37°C for 3 h.

*β*-galactosidase expression reactions was prepared as described above with 5 nM of DNA coding for *β*-galactosidase and incubated at 37°C for 3 h. The reaction was then diluted 50x in PBS and 2 *µ*L of the diluted solution was mixed with 20 *µ*L of chlorophenol red-*β*-D-galactopyranoside (1 mg/mL, Sigma) and measured (absorbance: 580 nm) on a SynergyMX platereader (BioTek).

Trehalase expression reaction was prepared as described above with 5 nM of DNA coding for trehalase and incubated at 37°C for 3 h, 2.5 *µ*L of the PURE reaction was mixed with 2.5 *µ*L trehalose (500mM) and incubated for 15 min at room temperature. After incubation, 5 *µ*L of DNS reagent was added, the final solution was incubated for 10 min at 99°C and 5 *µ*L was measured (absorbance: 540 nm) on a SynergyMX platereader (BioTek). DNS reagent was prepared by dissolving 5 mg of dinitrosalicylic acid (Acros Organics) in 250 *µ*L of water at 80°C. When this solution reaches room temperature, 100 *µ*L of NaOH, 2 N (Sigma) and 150 mg of potassium sodium tartrate-4-hydrate (Merck) were added and the volume is brought to a volume of 500 *µ*L with distilled water.

Repression of deGFP expression with zinc-finger transcription factors was measured in the reaction set-up as described above and supplemented with 800 nM of *E. coli* core RNAP and 4 *µ*M of *σ*70 factor, which were prepared as described previously (*16*). 1 nM of linear DNA template coding for deGPF and 1 nM of linear DNA template coding for ADD or CBD zinc-finger were used. The reaction was incubated at 37°C at constant shaking for 3 h, and measured (excitation: 488 nm, emission: 507 nm) on a SynergyMX platereader (BioTek).

### Cost calculations

To estimate the cost of PURE systems, we analyzed in detail the costs of the different subsets: protein components, ribosomes, and energy solution. The calculation for protein subset costs varies with the type of the system. For the TraMOS system, the reported cost of 0.052 USD/*µ*L was used (*17*). For our OnePot system, the cost was estimated based on the calculations given in Supplementary Table S1, with the assumptions that some of the materials can be reused and that four purifications can be done simultaneously in one working day. In the case of the HomeMade PURE system (Supplementary Table S6), our estimate was based on the price charged by the EPFL protein expression core facility: 300 USD per 2 L expression culture, which corresponds to our calculation for OnePot PURE of 83 USD per 0.5 L culture (332 USD for 2 L, Supplementary Table S1). Although the total price of this PURE system is high, the total amount of proteins purified is higher as well which can generate at least 40 mL of PURE HomeMade system (based on the volume of the protein limiting the preparation, in our case EF-Tu). Therefore, the price per *µ*L of HomeMade protein components is 0.27 USD.

Two different possibilities were taken into account in the case of the ribosome subset. In the first system, commercial ribosomes (Supplementary Table S7) were used for the PURE reactions (TraMOS). In the second system, purified ribosomes were used (HomeMade and OnePot PURE). The cost calculations for purified ribosomes are given in Supplementary Table S2, with the assumptions that some of the materials can be reused and that hands-on time for one purification is a single working day.

The cost calculation for the OnePot energy solution is described in Supplementary Table S3, with the assumption that half a day is necessary for the preparation of 20 mL of energy solution. For the TraMOS energy solution and the additional protein components, the costs were recalculated based on the component’s price that would apply for the preparation of the given solutions (Supplementary Table S7). For some of the additional protein components, we were not able to determine the exact protein which was purchased and its amount used, mostly due to a difference in the type of units reported in the paper as compared to the units specified by the supplier. However, we arrived at a very similar cost estimate as given in the original calculation. Furthermore, we assumed that the work required for the solution preparation is taken into account in the purification cost calculation, so we did not consider it.

In the case of PURExpress, the total cost was based on the commercial price. The values used in the cost calculation were derived from experience with the actual experiments while preparing the different subsets. All costs for the different components were based on the prices given in our internal EPFL system when performing the calculation; no delivery costs were taken into account.

### Important details and tips

1. The optimal concentration of *Mg*^2+^ in the energy solution is essential to high expression levels. If low expression levels are observed with an in-house prepared energy solution, we recommend to perform a *Mg*^2+^ titration.
2. tRNAs should not be weighed, but should be diluted directly in the flask, to avoid RNAase contamination (*30*).
3. All buffers should be sterile filtered to avoid bacterial contamination.
4. 2-mercaptoethanol should be added to the solutions immediately before use, buffers without 2-mercaptoethanol can be stored for an extended period in the fridge.
5. The overnight cultures should be shaken at 260 rpm, and well mixed prior to culture inoculation.
6. The expression cultures should be performed in a baffled flask to ensure proper oxygenation.

## Supporting information

## Data and material availability

All data presented is available upon request from the authors. Plasmids pET-21a(+)-IPP1-His and pET-21a(+)-AK1-His are available from AddGene.

## Supporting Information Available

The Supporting Information is available free of charge on the ACS Publications website.

Supporting Figures S1 - S12. Supporting Tables S1 - S12. (pdf)

Supporting Tables for LC-MS/MS data. (Excel)

### ORCID

Barbora Lavickova: 0000-0003-2903-2666

Sebastian J. Maerkl: 0000-0003-1917-5268

## Author Contributions

B.L. performed experiments. B.L. and S.J.M. designed experiments, analyzed data, and wrote the manuscript.

## Acknowledgement

We thank Yoshihiro Shimizu (RIKEN) for kindly supplying plasmids for the PURE system, members of the EPFL Protein Crystallography and Protein Expression Core Facilities, especially Jean Philippe Gaudry for providing us with purified PURE protein components and help with ribosome purification, EPFL ISICs Mass Spectrometry facility for helping us with sample analysis by mass spectrometry, Nadanai Laohakunakorn and Ekaterina Petrova for help with zinc-fingers transcription factors and trehalase assay, respectively. We also thank Corinna Tuckey and John DeMartino from NEB for providing us with custom PURExpress kits. This work was supported by the École Polytechnique Fédérale de Lausanne.

## Supporting Information

**Figure S1:**
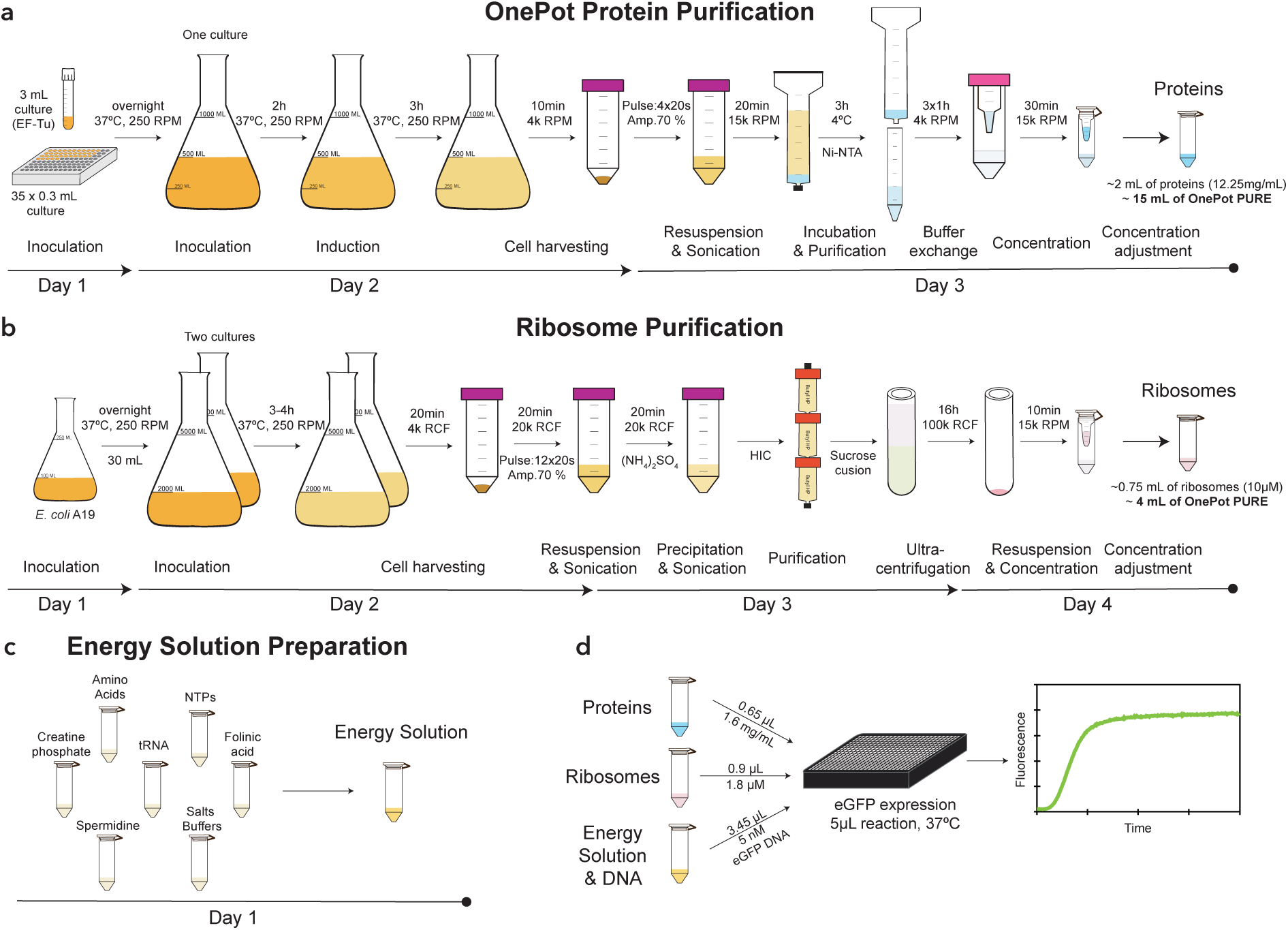
Schematics depicting all steps of the OnePot PURE production. **(a)** Protein purification, **(b)** ribosome purification, and **(c)** energy solution preparation steps. The description of the different steps as well as the day on which they are performed are indicated below the schematics. **(d)** Composition of the OnePot PURE reaction. Two numbers are given for each subset, the volume required for a 5 *µ*L reaction and the component concentration in the reaction.

**Figure S2:**
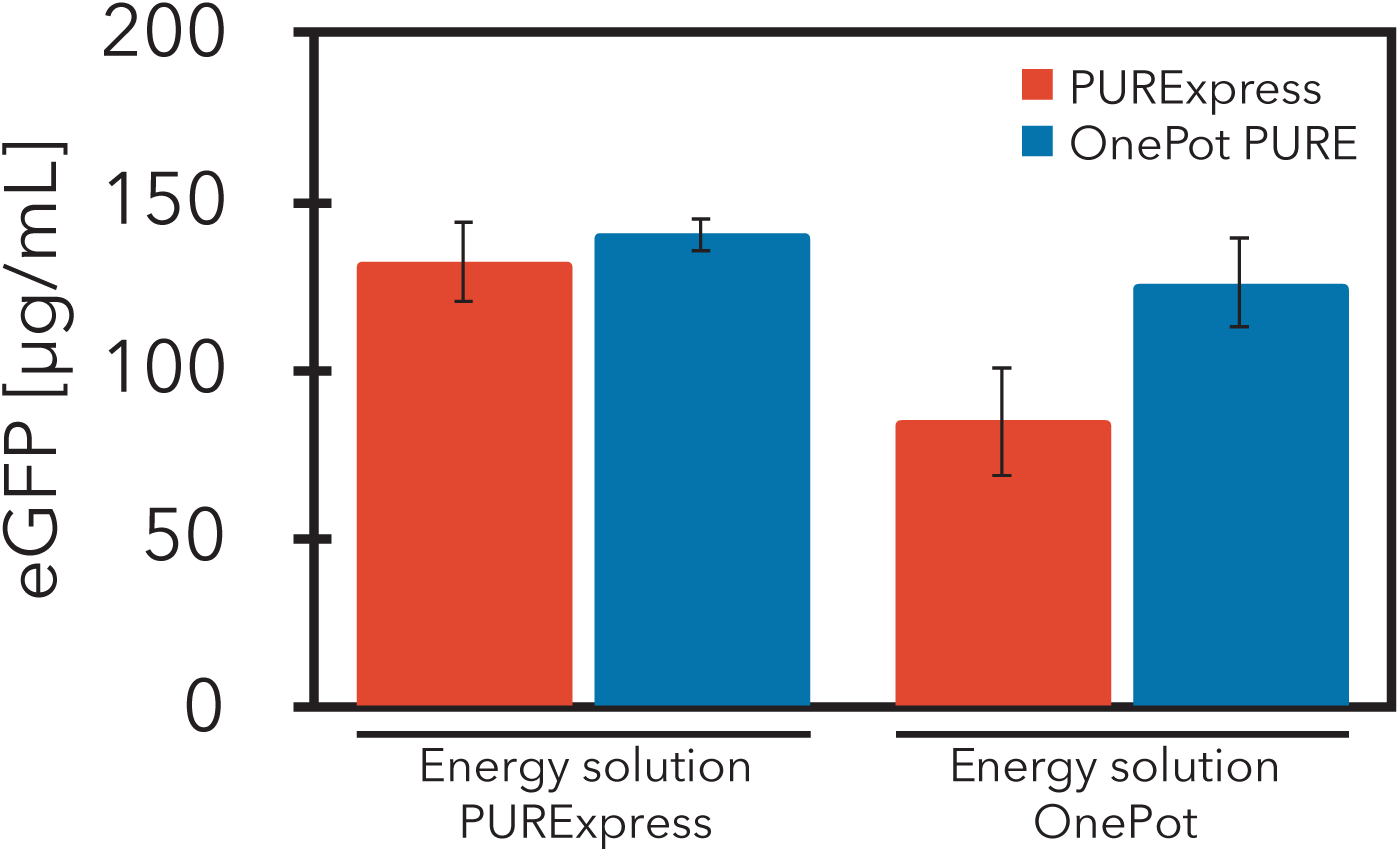
Comparison of eGFP expression levels in PURExpress (Solution B) and OnePot PURE (EF-Tu 47%, replicate A) supplied with commercial energy solution (Solution A, PURExpress) and the OnePot energy solution used in this study. Each data point represents at least five technical replicates (mean ± s.d.)

**Figure S3:**
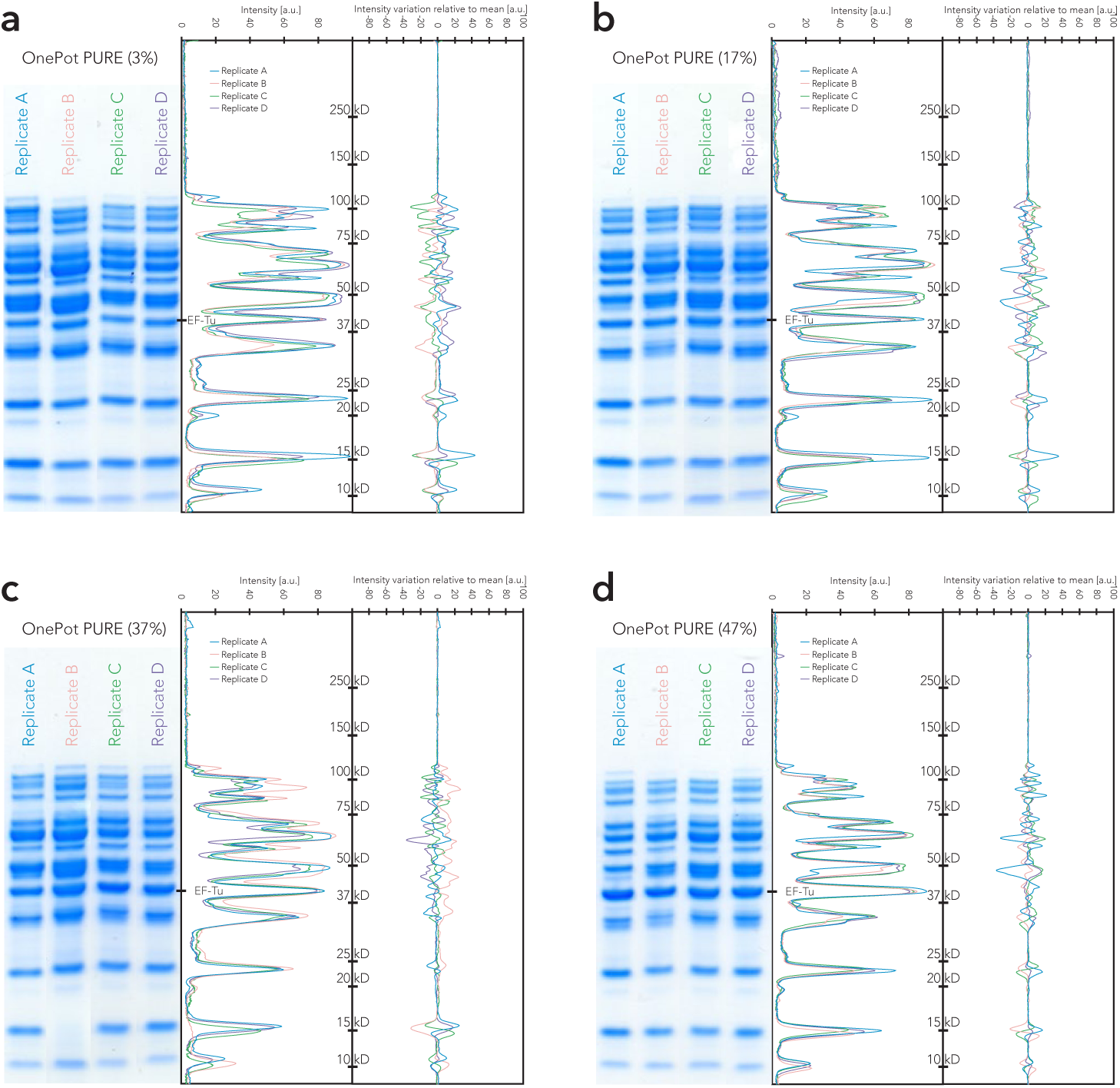
Coomassie blue stained SDS-PAGE gels of the four OnePot PURE formulations. **(a)** 3% EF-Tu, **(b)** 17% EF-Tu, **(c)** 37% EF-Tu, and **(d)** 47% EF-Tu. In the panels to the left of the gels, intensities of the different replicates are plotted with molecular weight standards (kDa). On the right the intensity variations relative to the inter-replicate mean is shown.

**Figure S4:**
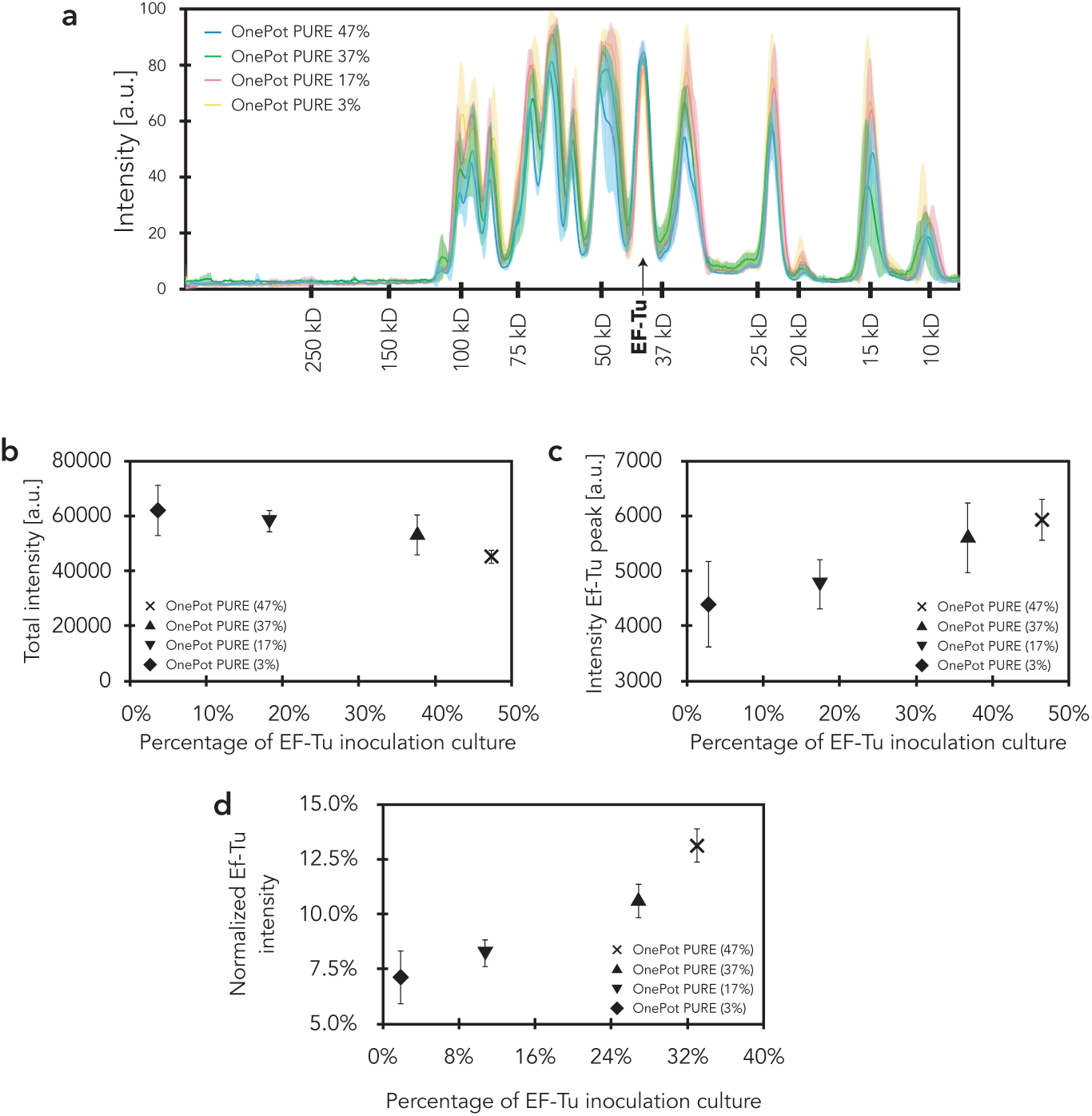
EF-Tu analysis. **(a)** Mean intensities of the different OnePot systems are plotted against molecular weight standards (kDa); the shaded regions represent the s.d. of the four biological replicates. **(b)** Total intensity of all protein bands as a function of EF-Tu clone inoculation percentage. **(c)** The integrated intensity of the EF-Tu peak from SDS-PAGE gel analysis as a function of EF-Tu clone inoculation percentage. **(d)** The normalised EF-Tu intensity (integrated EF-Tu peak intensity / total protein intensity) as a function of EF-Tu clone inoculation percentage. **(b) - (d)** Each data point represents four biological replicates (mean ± s.d.)

**Figure S5:**
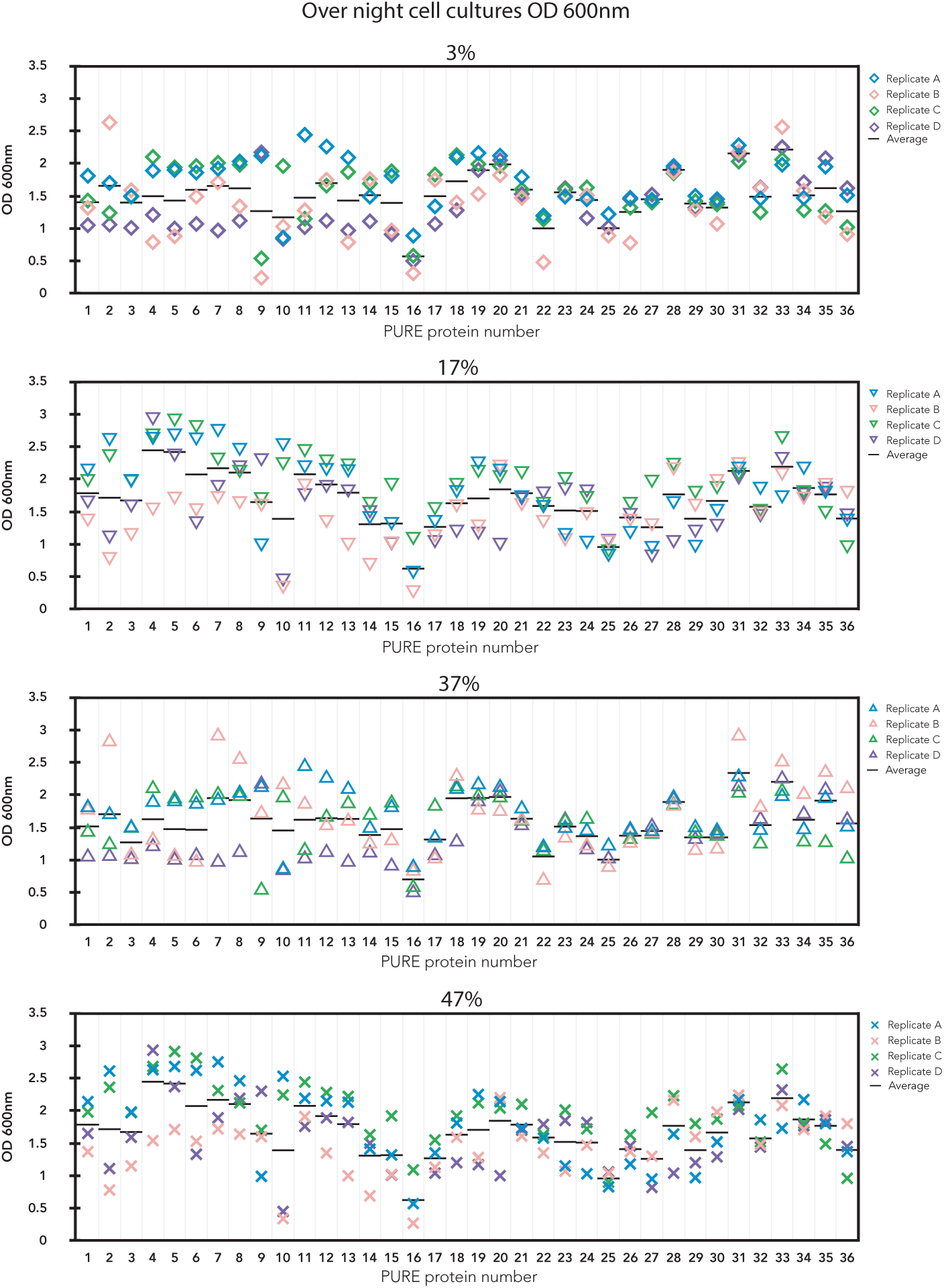
OD600 measurement of overnight cultures used for inoculation of the mixed culture.

**Figure S6:**
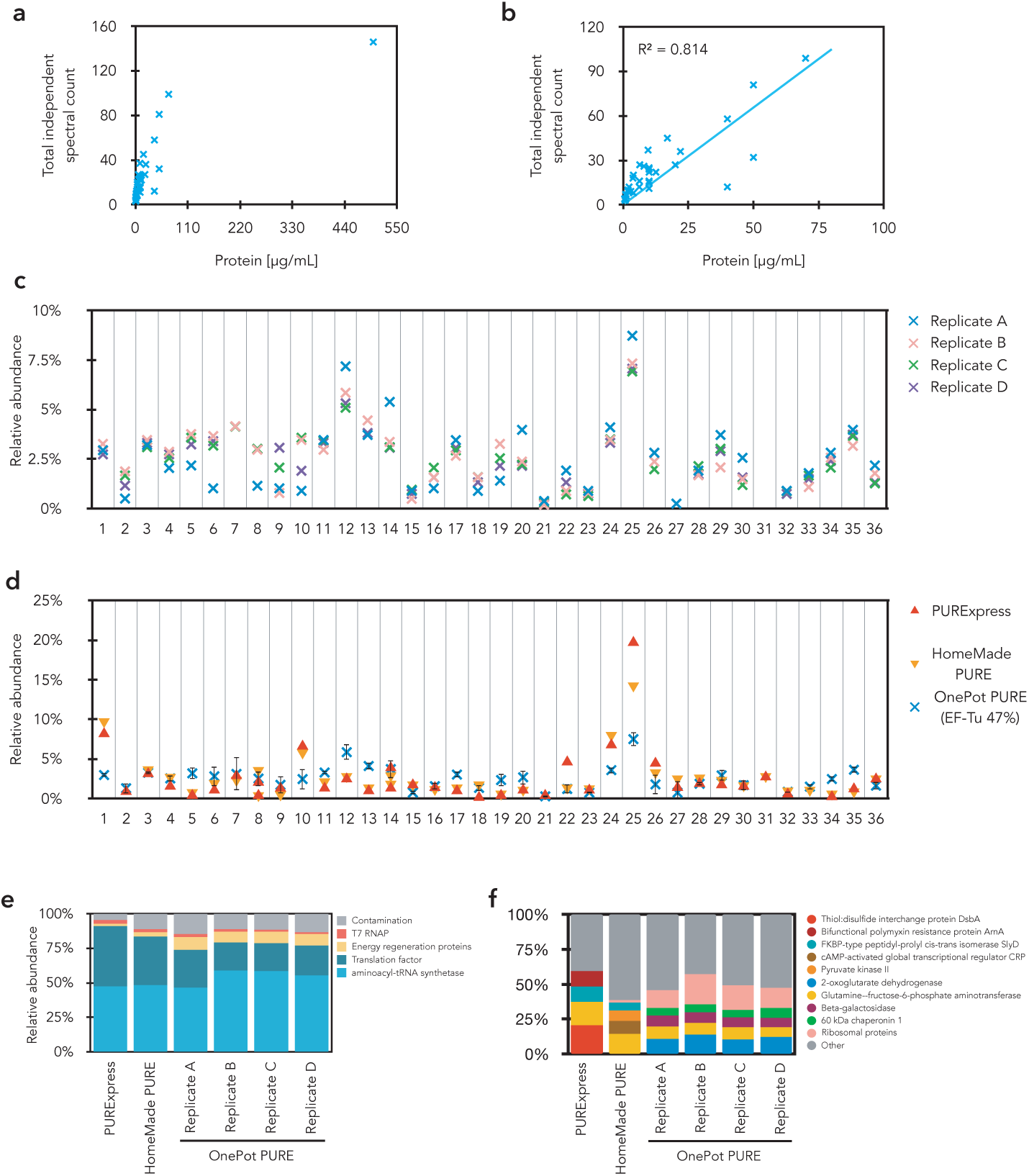
**(a)** Total independent spectral count from MS analysis of HomeMade PURE system vs protein concentration based on A280 in the HomeMade PURE system **(b)** Correlation of total independent spectral count and protein concentration (excluded EF-Tu). **(c)** Relative abundance of OnePot (EF-Tu 47%) system components normalized to total protein content based on total independent spectral count. **(d)** Relative abundance of PUR-Express, HomeMade PURE, OnePot (EF-Tu 47%) system components normalized to total protein content based on total independent spectral count. **(e)** Normalized composition of different PURE systems based on total independent spectral count. **(f)** Detailed description of contamination in the different PURE systems. Relative abundance of the four highest contaminant and ribosomal proteins are shown.

**Figure S7:**
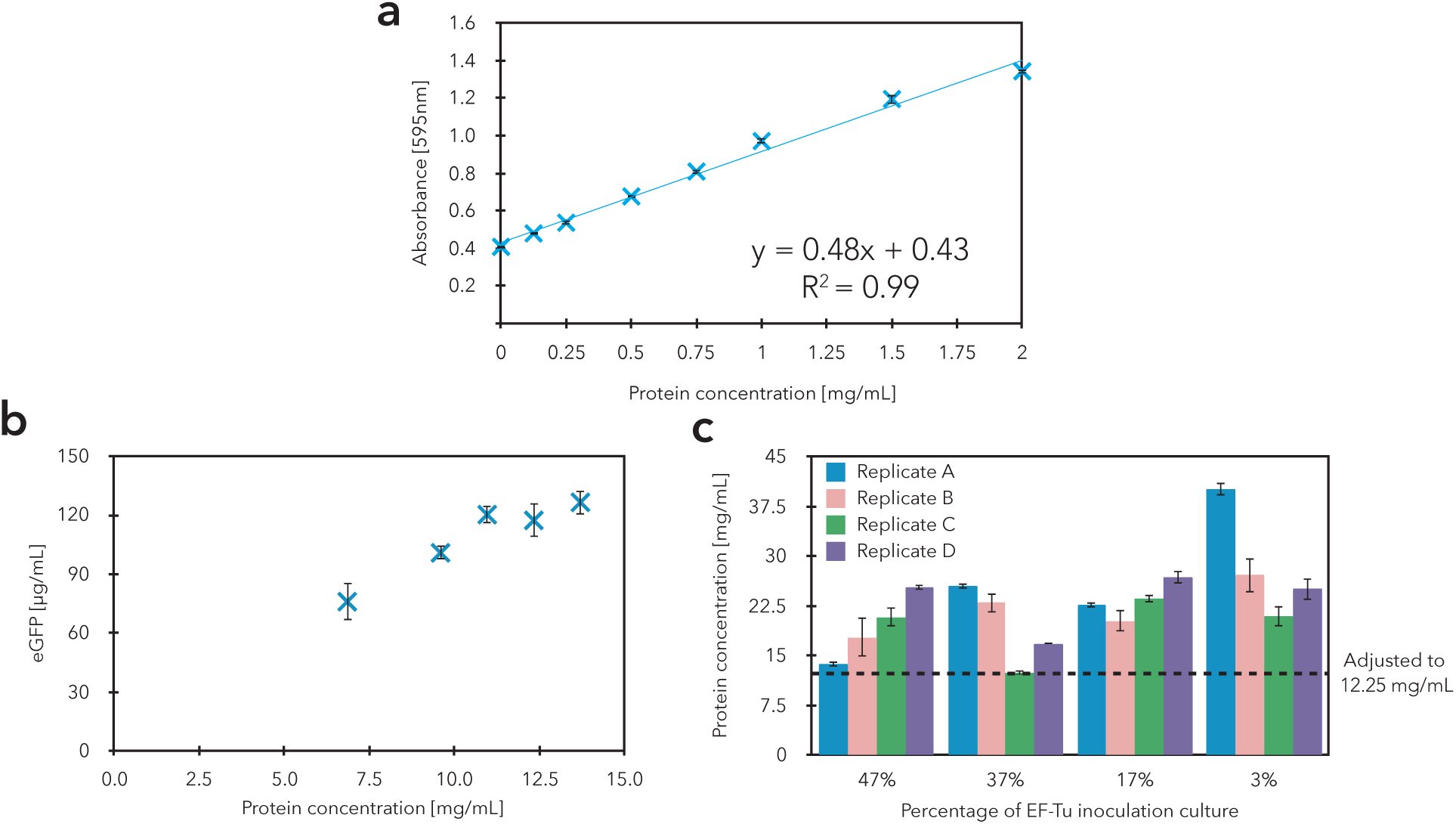
Protein concentration calibrations and adjustments. **(a)** Bradford assay standard calibration curve for protein concentration. The standard curve was produced by measuring the absorbance at 595 nm of prediluted bovine *γ*-globulin standards. Data are shown as mean ± s.d. (n = 3). Linear fit errors were not propagated as they were negligible compared to experimental errors. **(b)** eGFP expression as a function of protein concentrations in the protein subset of OnePot PURE (47%) replicate A (7.7× concentration in the final reaction). Each point represents at least two replicates; data are shown as mean ± s.d. **(c)** The concentrations of all OnePot protein subsets and their replicates after purification. Each bar represents two independent measurements in technical duplicate. Data are shown as mean ± s.d. The dotted line represents concentration (12.25 mg/mL, which is equal to 1.6 mg/mL in the final PURE reaction) to which all reactions were adjusted to.

**Figure S8:**
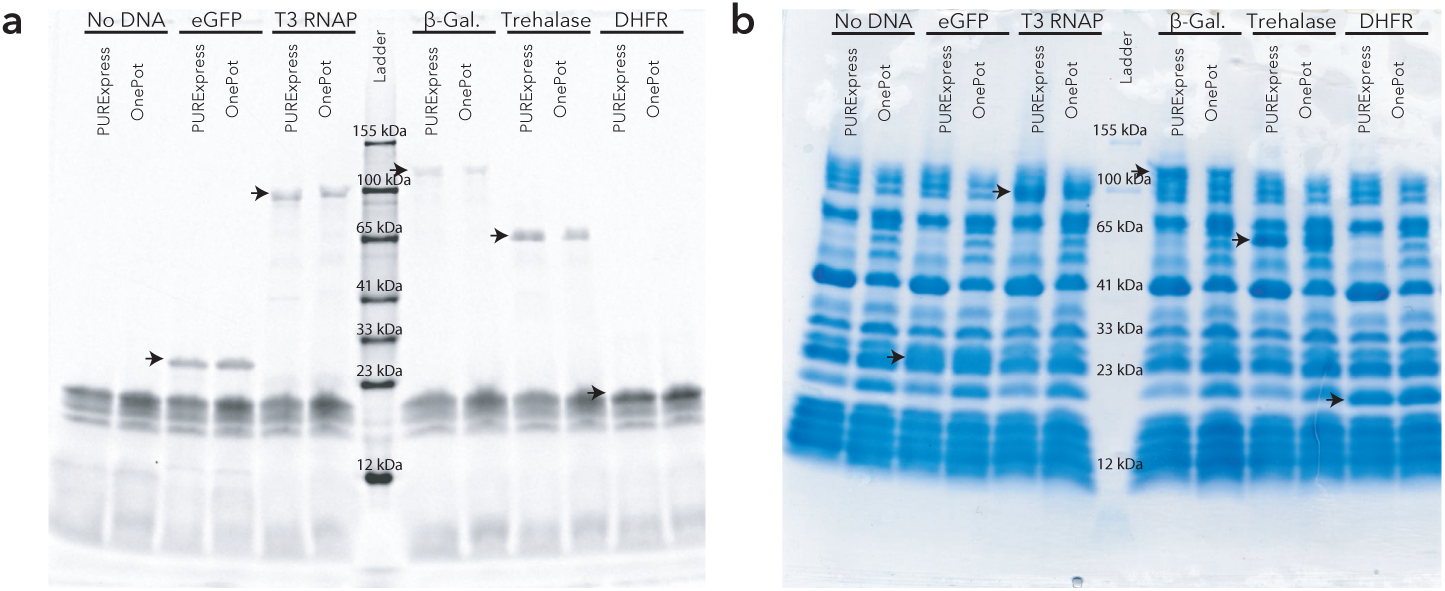
SDS-PAGE gel of proteins synthesized in PURExpress or OnePot (EF-Tu 47%, replicate A) **(a)** labeled with FluoroTect GreenLys, **(b)** Coomassie blue stained. Black arrows indicate the expected bands of synthesized proteins, GFP (26.9 kDa) T3 RNAP (98.8 kDa), *β*-galactosidase (116.5 kDa) and trehalase (63.7 kDa), DHFR (18 kDa)

**Figure S9:**
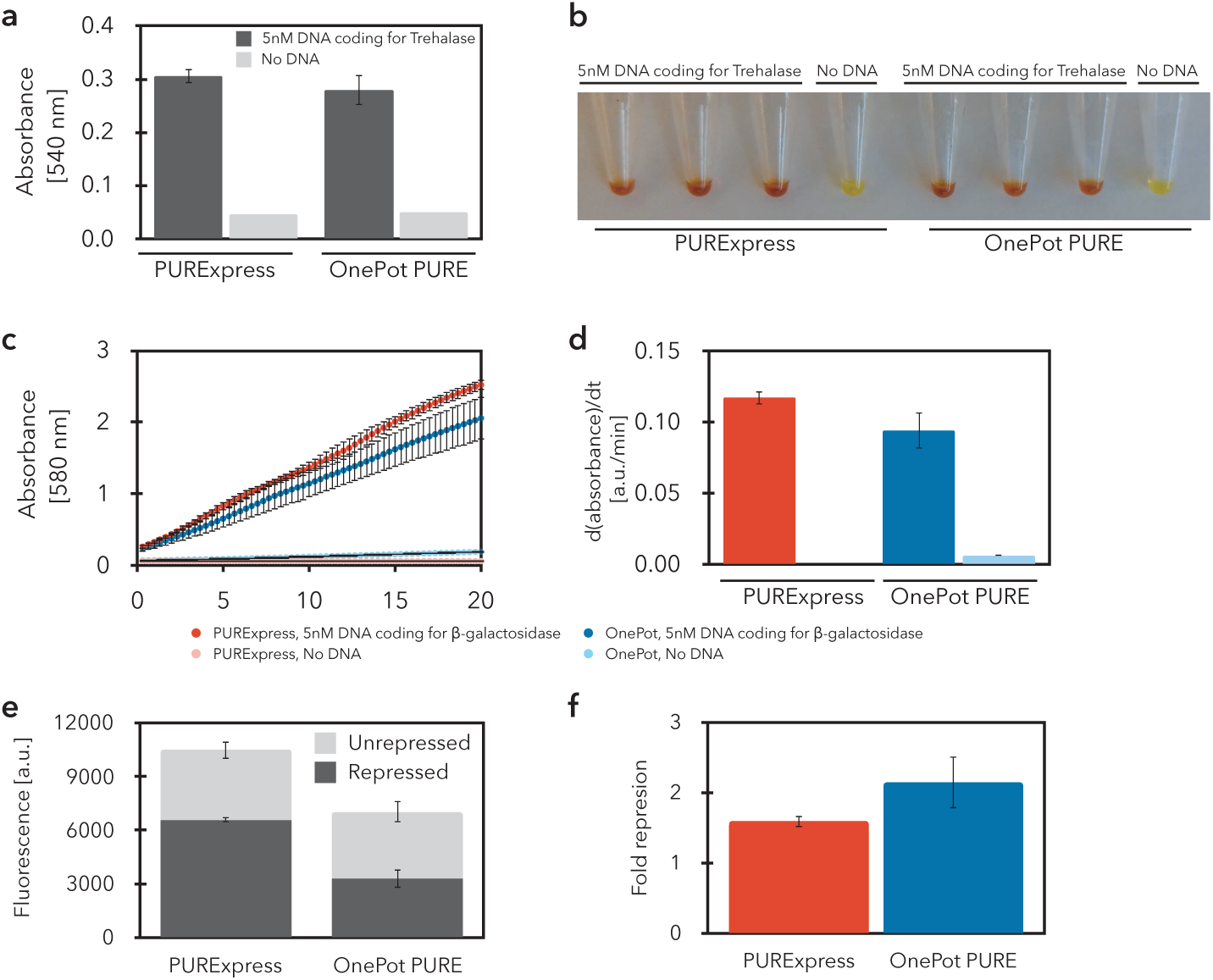
Activities of different proteins, expressed in PURExpress and OnePot (EF-Tu 47%, replicate A). Trehalase assay: **(a)** Absorbance change at 540 nm and **(b)** image of resulting color change due to the presence of trehalase in the reaction. Three reactions were measured for positive samples. Error bars represent standard deviation. *β*-galactosidase assay: **(c)** absorbance (580 nm) increase over time due to substrate cleavage, **(d)** slope of absorbance. Three and one OnePot PURE reactions were measured for positive and negative samples, respectively. Each reaction was measured in triplicate. Error bars represent standard deviation. Zinc-finger (ZF) repression: **(e)** Down-regulation of deGFP expression, due to binding of ZF to the target promoter. deGFP containing lambda PR promoter containing double ADD ZF binding sites was used as a reporter. The ADD ZF was co-expressed with deGFP (repressed state), and the CBD ZF was co-expressed as a negative control (unrepressed state). **(f)** Fold-repression, the ratio of unrepressed to repressed expression levels. Each data point represents three technical replicates (mean ± s.d.)

**Figure S10:**
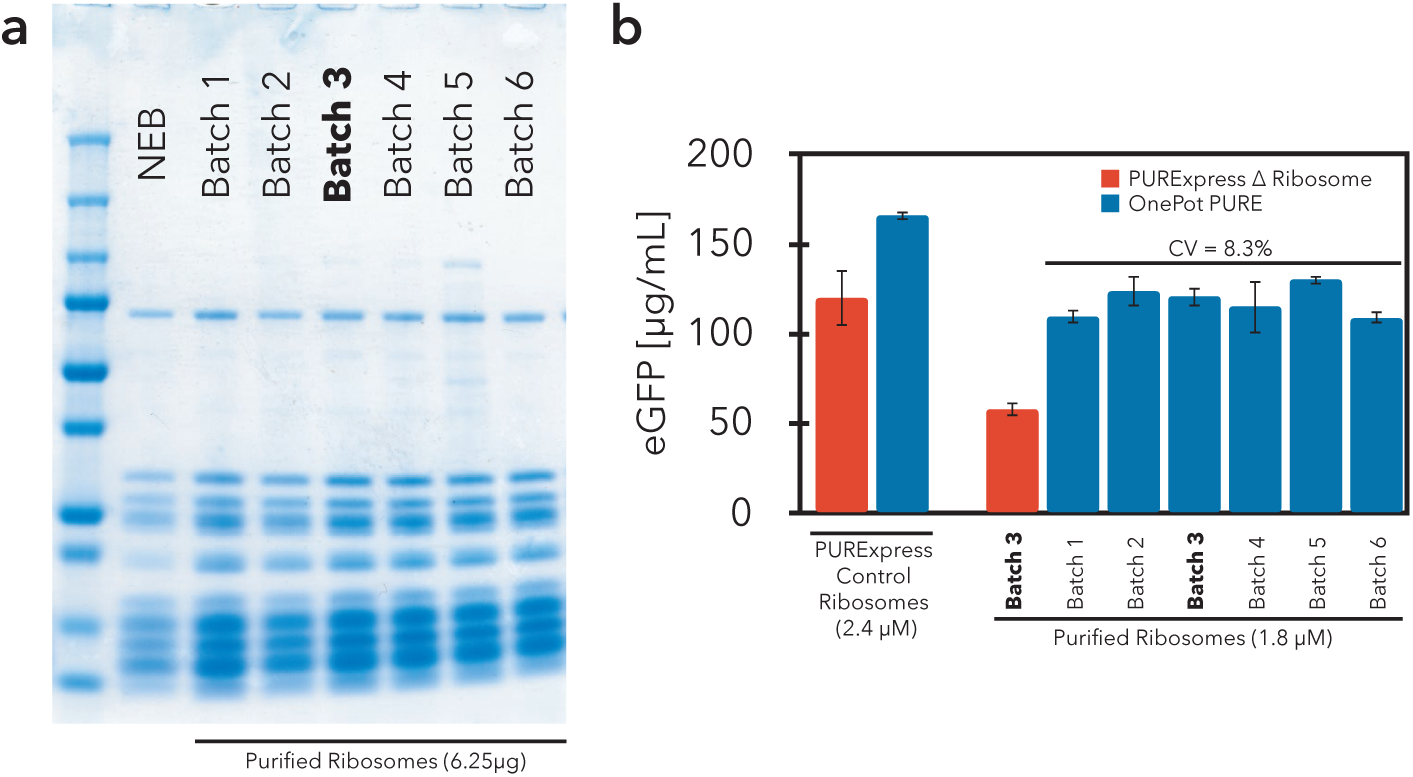
Comparison of commercial ribosomes (ribosomes from PURExpress ? ribosome kit, NEB) and different batches of ribosomes purified in our laboratory. Batch 3 was used throughout this study. **(a)** Coomassie blue stained SDS-PAGE gels of different ribosomes. The amounts loaded onto the gel were 6.24 *µ*g for NEB ribosomes and 6.25 *µ*g in the case of purified ribosomes. **(b)** Comparison of expression levels in PURExpress ? ribosome kit and OnePot PURE (EF-Tu 47%, replicate A) supplied with PURExpress control ribosomes (2.4 *µ*M) and purified ribosomes (1.8 *µ*M). Each data point represents two technical replicates (mean ± s.d.)

**Figure S11:**
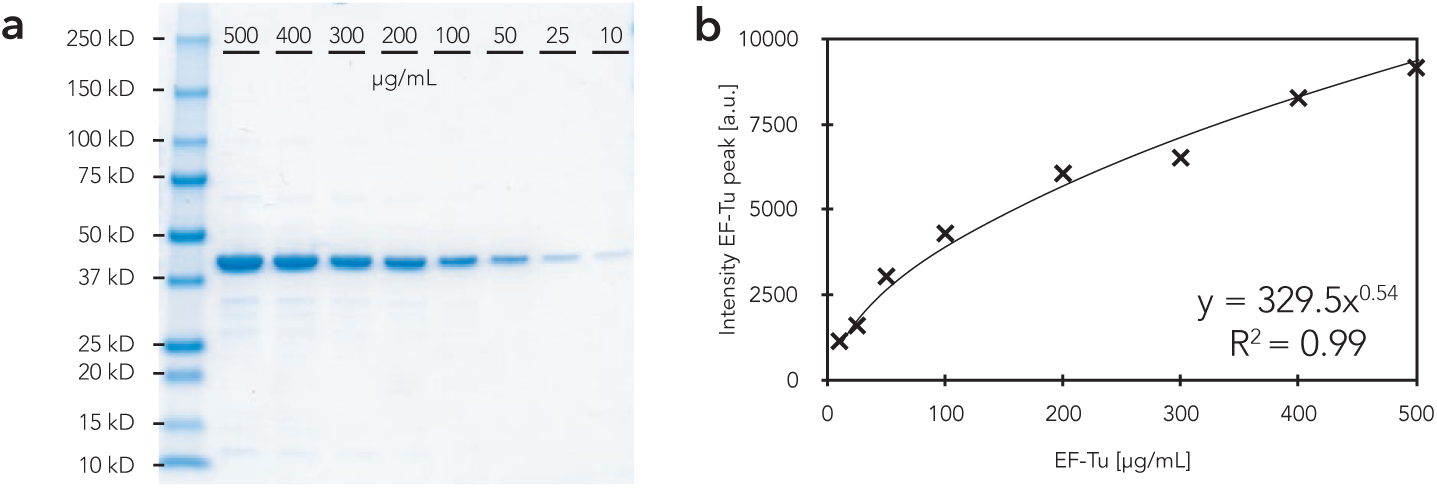
Standard calibration curve for EF-Tu protein concentration. **(a)** Coomassie blue stained SDS-PAGE gels of different concentrations of EF-Tu. **(b)** The standard curve was produced by measuring the integrated intensity of the EF-Tu peak at different EF-Tu concentrations. The reference EF-Tu concentration was determined by absorbance measurement at 280 nm.

**Figure S12:**
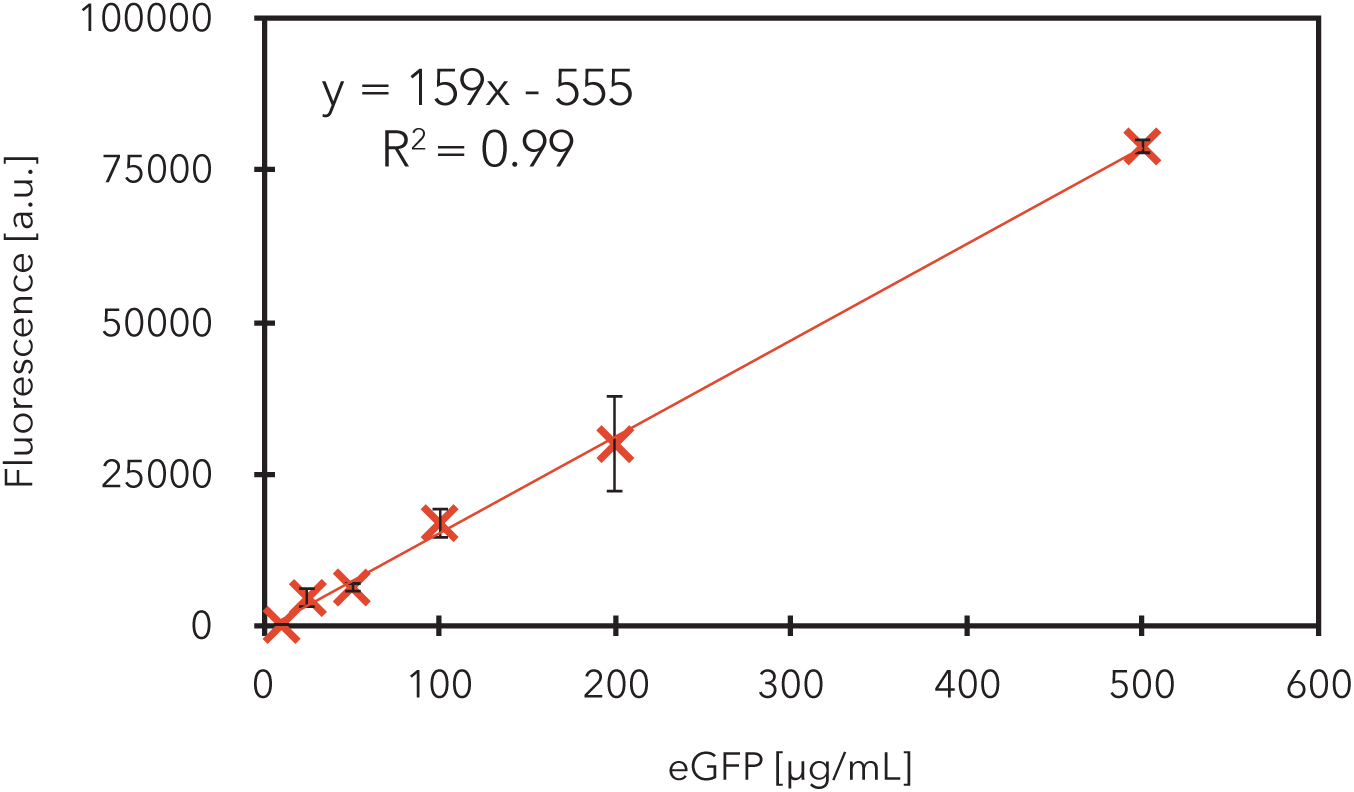
Standard calibration curve for eGFP. The standard curve was produced by measuring the fluorescence over 60 min for different eGFP (TP790050, AMS Biotechnology) concentrations in PBS on a plate reader with the same settings as for *in vitro* expression. Excitation and emission wavelengths were 488 nm and 507 nm, respectively. Experiments were performed in triplicates. Fluorescence measurements for the first 20 min were not considered. Data are shown as mean ± s.d. (n = 3). Linear fit errors were not propagated as they were negligible compared to experimental errors.

**Table S1:**
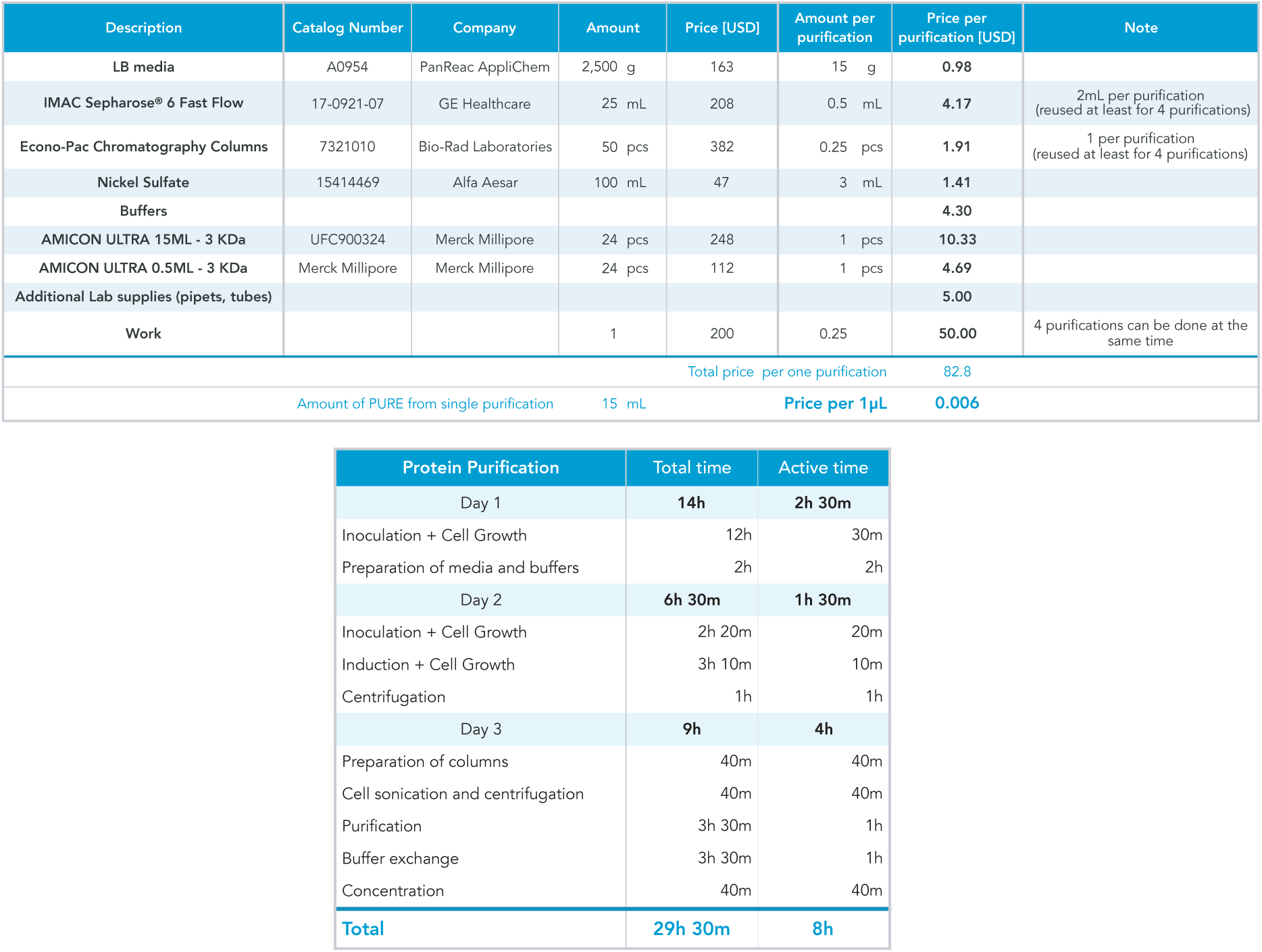
OnePot protocol cost and time estimate

**Table S2:** Ribosome protocol cost and time estimate

Ribosomes Purification
| Description | Catalog Number | Company | Amount | Price | Amount per purification | Price per purification [USD] | Note |
| --- | --- | --- | --- | --- | --- | --- | --- |
| LB media | A0954 | PanReac AppliChem | 2500 g | CHF163.27 | 100 g | 6.53 |  |
| HiTrap Butyl HP Column | 28411005 | GE Healthcare | 5 pcs | CHF322.73 | 0.2 pcs | 12.91 | 2-3 per purification (reused for multiple purifications) |
| Thickwall Polycarbonate Tube | 355631 | Beckman | 25 pcs | CHF274.63 | 1 pcs | 10.99 | reused for multiple purifications |
| Whatman® GD/X syringe filters | WHA68722504 | GE Whatman | 50 pcs | CHF303.68 | 1 pcs | 6.07 |  |
| Buffers |  |  |  |  |  | 22.63 |  |
| AMICON ULTRA 0.5ML - 3 KDa | Merck Millipore | Merck Millipore | 24 pcs | CHF112.45 | 1 pcs | 4.69 |  |
| Additional Lab supplies (pipets, tubes) |  |  |  |  |  | 5.00 |  |
| Work |  |  | 1 | CHF200.00 | 1 | 200.00 |  |
| Total price per one purification |  |  |  |  |  | 268.8 |  |
| Amount of PURE from single purification |  |  | 4 mL | Price per 1µL |  | 0.07 |  |

| Ribosome Purification | Total time | Active time |
| --- | --- | --- |
| Day 1 | 14h | 2h 10m |
| Inoculation + Cell Growth | 12h | 10m |
| Preparation of media and buffers | 2h | 2h |
| Day 2 | 5h | 1h 10m |
| Inoculation + Cell Growth | 4h | 10m |
| Centrifugation | 1h | 1h |
| Day 3 | 21h | 3h 40m |
| Preparation and cleaning of columns | 2h | 1h |
| Cell sonication and centrifugation | 1h 30m | 40m |
| Purification | 1h 30m | 1h 30m |
| Ultracentrifugation | 16h | 30m |
| Day 3 | 1h | 1h |
| Resuspension, Concentration | 1h | 1h |
| **Total** | **41h** | **8h** |

**Table S3:**
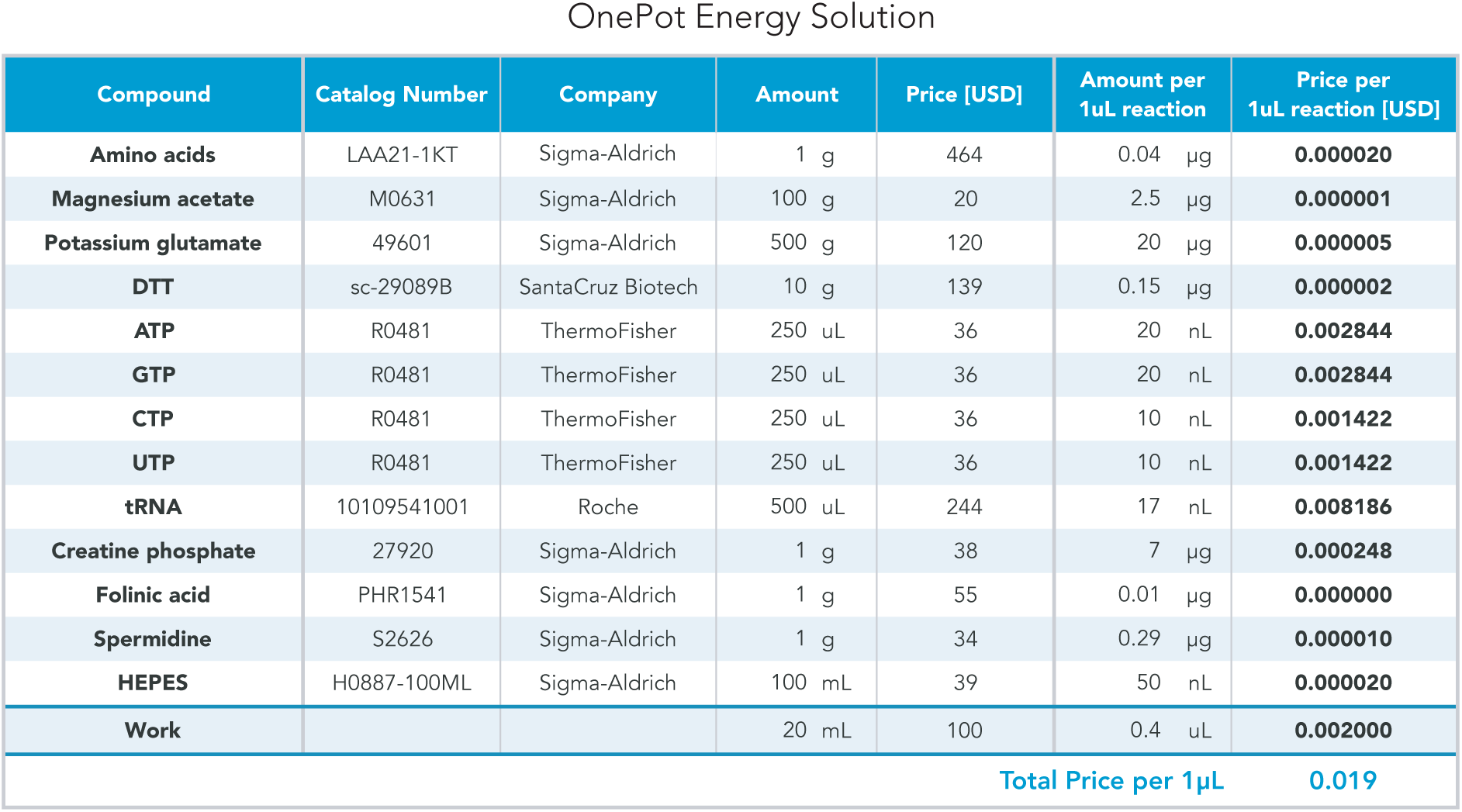
Energy solution cost estimates

OnePot Energy Solution
| Compound | Catalog Number | Company | Amount | Price [USD] | Amount per 1uL reaction | Price per 1uL reaction [USD] |
| --- | --- | --- | --- | --- | --- | --- |
| **Amino acids** | LAA21-1KT | Sigma-Aldrich | 1 g | 464 | 0.04 µg | **0.000020** |
| **Magnesium acetate** | M0631 | Sigma-Aldrich | 100 g | 20 | 2.5 µg | **0.000001** |
| **Potassium glutamate** | 49601 | Sigma-Aldrich | 500 g | 120 | 20 µg | **0.000005** |
| **DTT** | sc-29089B | SantaCruz Biotech | 10 g | 139 | 0.15 µg | **0.000002** |
| **ATP** | R0481 | ThermoFisher | 250 uL | 36 | 20 nL | **0.002844** |
| **GTP** | R0481 | ThermoFisher | 250 uL | 36 | 20 nL | **0.002844** |
| **CTP** | R0481 | ThermoFisher | 250 uL | 36 | 10 nL | **0.001422** |
| **UTP** | R0481 | ThermoFisher | 250 uL | 36 | 10 nL | **0.001422** |
| **tRNA** | 10109541001 | Roche | 500 uL | 244 | 17 nL | **0.008186** |
| **Creatine phosphate** | 27920 | Sigma-Aldrich | 1 g | 38 | 7 µg | **0.000248** |
| **Folinic acid** | PHR1541 | Sigma-Aldrich | 1 g | 55 | 0.01 µg | **0.000000** |
| **Spermidine** | S2626 | Sigma-Aldrich | 1 g | 34 | 0.29 µg | **0.000010** |
| **HEPES** | H0887-100ML | Sigma-Aldrich | 100 mL | 39 | 50 nL | **0.000020** |
| **Work** |  |  | 20 mL | 100 | 0.4 uL | **0.002000** |
| Total Price per 1µL |  |  |  |  |  | **0.019** |

**Table S4:** OnePot inoculation culture volumes

| Number | Protein | Vector | Strain | OnePot (3%) |  | OnePot (17%) |  | OnePot (37%) |  | OnePot (47%) |  |
| --- | --- | --- | --- | --- | --- | --- | --- | --- | --- | --- | --- |
| Amount of inoculation culture |  |  |  | μL | % | μL | % | μL | % | μL | % |
| 1 | AlaRS | pQE30 | M15 | 100 | 2.8% | 85 | 2.4% | 65 | 1.8% | 55 | 1.5% |
| 2 | ArgRS | pET16b | BL21(DE3) | 100 | 2.8% | 85 | 2.4% | 65 | 1.8% | 55 | 1.5% |
| 3 | AsnRS | pQE30 | M15 | 100 | 2.8% | 85 | 2.4% | 65 | 1.8% | 55 | 1.5% |
| 4 | AspRS | pET21a | BL21(DE3) | 100 | 2.8% | 85 | 2.4% | 65 | 1.8% | 55 | 1.5% |
| 5 | CysRS | pET21a | BL21(DE3) | 100 | 2.8% | 85 | 2.4% | 65 | 1.8% | 55 | 1.5% |
| 6 | GlnRS | pET21a | BL21(DE3) | 100 | 2.8% | 85 | 2.4% | 65 | 1.8% | 55 | 1.5% |
| 7 | GluRS | pET21a | BL21(DE3) | 100 | 2.8% | 85 | 2.4% | 65 | 1.8% | 55 | 1.5% |
| 8 | GlyRS | pET21a | BL21(DE3) | 100 | 2.8% | 85 | 2.4% | 65 | 1.8% | 55 | 1.5% |
| 9 | HisRS | pET21a | BL21(DE3) | 100 | 2.8% | 85 | 2.4% | 65 | 1.8% | 55 | 1.5% |
| 10 | IleRS | pET21a | BL21(DE3) | 100 | 2.8% | 85 | 2.4% | 65 | 1.8% | 55 | 1.5% |
| 11 | LeuRS | pET21a | BL21(DE3) | 100 | 2.8% | 85 | 2.4% | 65 | 1.8% | 55 | 1.5% |
| 12 | LysRS | pET21a | BL21(DE3) | 100 | 2.8% | 85 | 2.4% | 65 | 1.8% | 55 | 1.5% |
| 13 | MetRS | pET21a | BL21(DE3) | 100 | 2.8% | 85 | 2.4% | 65 | 1.8% | 55 | 1.5% |
| 14 | PheRS | pQE30 | M15 | 100 | 2.8% | 85 | 2.4% | 65 | 1.8% | 55 | 1.5% |
| 15 | ProRS | pET21a | BL21(DE3) | 100 | 2.8% | 85 | 2.4% | 65 | 1.8% | 55 | 1.5% |
| 16 | SerRS | pET21a | BL21(DE3) | 100 | 2.8% | 85 | 2.4% | 65 | 1.8% | 55 | 1.5% |
| 17 | ThrRS | pQE30 | M15 | 100 | 2.8% | 85 | 2.4% | 65 | 1.8% | 55 | 1.5% |
| 18 | TrpRS | pET21a | BL21(DE3) | 100 | 2.8% | 85 | 2.4% | 65 | 1.8% | 55 | 1.5% |
| 19 | TyrRS | pET21a | BL21(DE3) | 100 | 2.8% | 85 | 2.4% | 65 | 1.8% | 55 | 1.5% |
| 20 | ValRS | pET21a | BL21(DE3) | 100 | 2.8% | 85 | 2.4% | 65 | 1.8% | 55 | 1.5% |
| 21 | IF1 | pQE30 | M15 | 100 | 2.8% | 85 | 2.4% | 65 | 1.8% | 55 | 1.5% |
| 22 | IF2 | pQE30 | M15 | 100 | 2.8% | 85 | 2.4% | 65 | 1.8% | 55 | 1.5% |
| 23 | IF3 | pQE30 | M15 | 100 | 2.8% | 85 | 2.4% | 65 | 1.8% | 55 | 1.5% |
| 24 | EF-G | pQE60 | M15 | 100 | 2.8% | 85 | 2.4% | 65 | 1.8% | 55 | 1.5% |
| 25 | EF-Tu | pQE60 | M15 | 100 | 2.8% | 625 | 17.4% | 1325 | 36.8% | 1675 | 46.5% |
| 26 | EF-Ts | pQE60 | M15 | 100 | 2.8% | 85 | 2.4% | 65 | 1.8% | 55 | 1.5% |
| 27 | RF1 | pQE30 | M15 | 100 | 2.8% | 85 | 2.4% | 65 | 1.8% | 55 | 1.5% |
| 28 | RF2 | pET15b | BL21(DE3) | 100 | 2.8% | 85 | 2.4% | 65 | 1.8% | 55 | 1.5% |
| 29 | RF3 | pQE30 | M15 | 100 | 2.8% | 85 | 2.4% | 65 | 1.8% | 55 | 1.5% |
| 30 | RRF | pQE60 | M15 | 100 | 2.8% | 85 | 2.4% | 65 | 1.8% | 55 | 1.5% |
| 31 | MTF | pET21a | BL21(DE3) | 100 | 2.8% | 85 | 2.4% | 65 | 1.8% | 55 | 1.5% |
| 32 | CK | pQE30 | M15 | 100 | 2.8% | 85 | 2.4% | 65 | 1.8% | 55 | 1.5% |
| 33 | MK | pET21a | BL21(DE3) | 100 | 2.8% | 85 | 2.4% | 65 | 1.8% | 55 | 1.5% |
| 34 | NDK | pQE30 | M15 | 100 | 2.8% | 85 | 2.4% | 65 | 1.8% | 55 | 1.5% |
| 35 | PPIase | pET21a | BL21(DE3) | 100 | 2.8% | 85 | 2.4% | 65 | 1.8% | 55 | 1.5% |
| 36 | T7 RNAP | pQE30 | M15 | 100 | 2.8% | 85 | 2.4% | 65 | 1.8% | 55 | 1.5% |
| Total amount of inoculation culture |  |  |  | 3600 | 100% | 3600 | 100% | 3600 | 100% | 3600 | 100% |

**Table S5:** HomeMade PURE protein concentrations

| Number | Protein | Vector | Antibiotic | Strain | HomeMade PURE |  |
| --- | --- | --- | --- | --- | --- | --- |
|  |  |  |  |  | Final Concentration in reaction [µg/mL] | Concentration in PURE protein solution [µg/mL] |
| 1 | AlaRS | pQE30 | Amp, Kan | M15 | 70 | 538 |
| 2 | ArgRS | pET16b | Amp | BL21(DE3) | 2 | 15 |
| 3 | AsnRS | pQE30 | Amp, Kan | M15 | 22 | 169 |
| 4 | AspRS | pET21a | Amp | BL21(DE3) | 8 | 62 |
| 5 | CysRS | pET21a | Amp | BL21(DE3) | 1 | 9 |
| 6 | GlnRS | pET21a | Amp | BL21(DE3) | 4 | 29 |
| 7 | GluRS | pET21a | Amp | BL21(DE3) | 13 | 97 |
| 8 | GlyRS | pET21a | Amp | BL21(DE3) | 10 | 74 |
| 9 | HisRS | pET21a | Amp | BL21(DE3) | 1 | 6 |
| 10 | IleRS | pET21a | Amp | BL21(DE3) | 40 | 308 |
| 11 | LeuRS | pET21a | Amp | BL21(DE3) | 4 | 31 |
| 12 | LysRS | pET21a | Amp | BL21(DE3) | 6 | 49 |
| 13 | MetRS | pET21a | Amp | BL21(DE3) | 2 | 18 |
| 14 | PheRS | pQE30 | Amp, Kan | M15 | 17 | 131 |
| 15 | ProRS | pET21a | Amp | BL21(DE3) | 10 | 77 |
| 16 | SerRS | pET21a | Amp | BL21(DE3) | 2 | 15 |
| 17 | ThrRS | pQE30 | Amp, Kan | M15 | 6 | 48 |
| 18 | TrpRS | pET21a | Amp | BL21(DE3) | 6 | 48 |
| 19 | TyrRS | pET21a | Amp | BL21(DE3) | 1 | 5 |
| 20 | ValRS | pET21a | Amp | BL21(DE3) | 2 | 14 |
| 21 | IF1 | pQE30 | Amp, Kan | M15 | 10 | 77 |
| 22 | IF2 | pQE30 | Amp, Kan | M15 | 40 | 308 |
| 23 | IF3 | pQE30 | Amp, Kan | M15 | 10 | 77 |
| 24 | EF-G | pQE60 | Amp, Kan | M15 | 50 | 385 |
| 25 | EF-Tu | pQE60 | Amp, Kan | M15 | 500 | 3846 |
| 26 | EF-Ts | pQE60 | Amp, Kan | M15 | 50 | 385 |
| 27 | RF1 | pQE30 | Amp, Kan | M15 | 10 | 77 |
| 28 | RF2 | pET15b | Amp | BL21(DE3) | 10 | 77 |
| 29 | RF3 | pQE30 | Amp, Kan | M15 | 10 | 77 |
| 30 | RRF | pQE60 | Amp, Kan | M15 | 10 | 77 |
| 31 | MTF | pET21a | Amp | BL21(DE3) | 20 | 154 |
| 32 | CK | pQE30 | Amp, Kan | M15 | 4 | 31 |
| 33 | MK | pET29b | Kan | BL21(DE3) | 3 | 23 |
| 34 | NDK | pQE30 | Amp, Kan | M15 | 1 | 8 |
| 35 | PPIase | pET29b | Kan | BL21(DE3) | 1 | 8 |
| 36 | T7 RNAP | pQE30 | Amp, Kan | M15 | 10 | 77 |
| Total protein concentration [µg/mL] |  |  |  |  | 966 | 7428 |

**Table S6:** HomeMade protocol cost estimates

| Description | Catalog Number | Amount per 1μL reaction | Amount per purification | Price per purification [USD] |
| --- | --- | --- | --- | --- |
| 1 | AlaRS | 70 ng | 49 mg | 300 |
| 2 | ArgRS | 2 ng | 6 mg | 300 |
| 3 | AsnRS | 8 ng | 93 mg | 300 |
| 4 | AspRS | 1 ng | 34 mg | 300 |
| 5 | CysRS | 4 ng | 48 mg | 300 |
| 6 | GlnRS | 10 ng | 38 mg | 300 |
| 7 | GluRS | 2 ng | 69 mg | 300 |
| 8 | GlyRS | 17 ng | 20 mg | 300 |
| 9 | HisRS | 10 ng | 85 mg | 300 |
| 10 | IleRS | 6 ng | 26 mg | 300 |
| 11 | LeuRS | 22 ng | 33 mg | 300 |
| 12 | LysRS | 13 ng | 112 mg | 300 |
| 13 | MetRS | 1 ng | 39 mg | 300 |
| 14 | PheRS | 40 ng | 27 mg | 300 |
| 15 | ProRS | 4 ng | 16 mg | 300 |
| 16 | SerRS | 6 ng | 42 mg | 300 |
| 17 | ThrRS | 2 ng | 67 mg | 300 |
| 18 | TrpRS | 6 ng | 64 mg | 300 |
| 19 | TyrRS | 1 ng | 56 mg | 300 |
| 20 | ValRS | 2 ng | 26 mg | 300 |
| 21 | IF1 | 10 ng | 14 mg | 300 |
| 22 | IF2 | 40 ng | 16 mg | 300 |
| 23 | IF3 | 10 ng | 36 mg | 300 |
| 24 | EF-G | 50 ng | 56 mg | 300 |
| 25 | EF-Tu | 500 ng | 20 mg | 300 |
| 26 | EF-Ts | 50 ng | 36 mg | 300 |
| 27 | RF1 | 10 ng | 37 mg | 300 |
| 28 | RF2 | 10 ng | 16 mg | 300 |
| 29 | RF3 | 10 ng | 66 mg | 300 |
| 30 | RRF | 10 ng | 8 mg | 300 |
| 31 | MTF | 20 ng | 36 mg | 300 |
| 32 | CK | 4 ng | 12 mg | 300 |
| 33 | MK | 3 ng | 41 mg | 300 |
| 34 | NDK | 1 ng | 119 mg | 300 |
| 35 | PPiase | 1 ng | 149 mg | 300 |
| 36 | T7 pol. | 10 ng | 20.8 mg | 300 |
| Total price per purifications |  |  |  | 10800 |
| Amount of PURE from single run of purifications (based on EF-Tu) | | 40 mL | Price per 1μL | 0.27 |

**Table S7:**
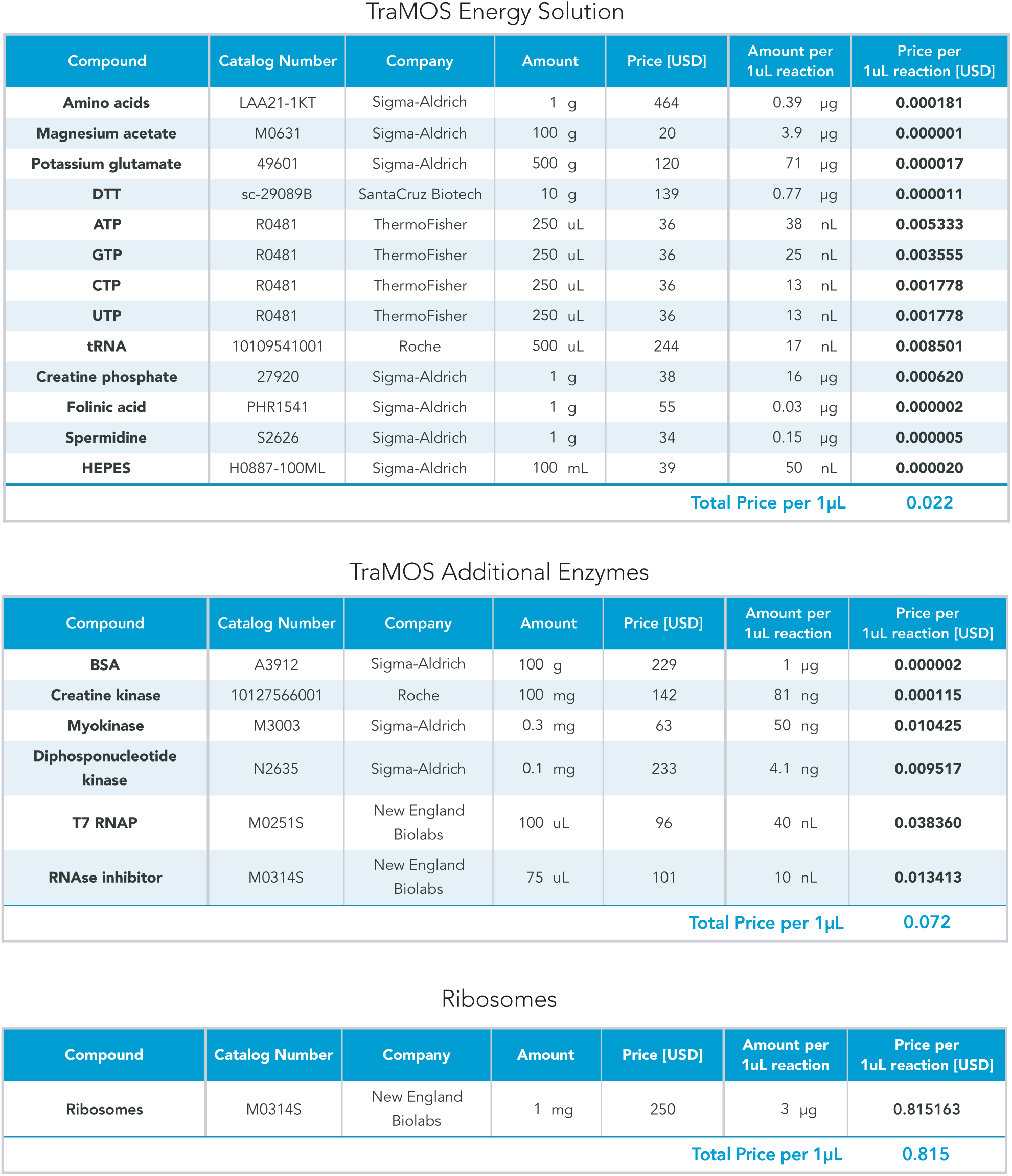
Energy solution and ribosome cost estimates for TraMOS

TraMOS Energy Solution
| Compound | Catalog Number | Company | Amount | Price [USD] | Amount per 1uL reaction | Price per 1uL reaction [USD] |
| --- | --- | --- | --- | --- | --- | --- |
| **Amino acids** | LAA21-1KT | Sigma-Aldrich | 1 g | 464 | 0.39 µg | **0.000181** |
| **Magnesium acetate** | M0631 | Sigma-Aldrich | 100 g | 20 | 3.9 µg | **0.000001** |
| **Potassium glutamate** | 49601 | Sigma-Aldrich | 500 g | 120 | 71 µg | **0.000017** |
| **DTT** | sc-29089B | SantaCruz Biotech | 10 g | 139 | 0.77 µg | **0.000011** |
| **ATP** | R0481 | ThermoFisher | 250 uL | 36 | 38 nL | **0.005333** |
| **GTP** | R0481 | ThermoFisher | 250 uL | 36 | 25 nL | **0.003555** |
| **CTP** | R0481 | ThermoFisher | 250 uL | 36 | 13 nL | **0.001778** |
| **UTP** | R0481 | ThermoFisher | 250 uL | 36 | 13 nL | **0.001778** |
| **tRNA** | 10109541001 | Roche | 500 uL | 244 | 17 nL | **0.008501** |
| **Creatine phosphate** | 27920 | Sigma-Aldrich | 1 g | 38 | 16 µg | **0.000620** |
| **Folinic acid** | PHR1541 | Sigma-Aldrich | 1 g | 55 | 0.03 µg | **0.000002** |
| **Spermidine** | S2626 | Sigma-Aldrich | 1 g | 34 | 0.15 µg | **0.000005** |
| **HEPES** | H0887-100ML | Sigma-Aldrich | 100 mL | 39 | 50 nL | **0.000020** |
| Total Price per 1µL |  |  |  |  |  | **0.022** |

TraMOS Additional Enzymes
| Compound | Catalog Number | Company | Amount | Price [USD] | Amount per 1uL reaction | Price per 1uL reaction [USD] |
| --- | --- | --- | --- | --- | --- | --- |
| **BSA** | A3912 | Sigma-Aldrich | 100 g | 229 | 1 µg | **0.000002** |
| **Creatine kinase** | 10127566001 | Roche | 100 mg | 142 | 81 ng | **0.000115** |
| **Myokinase** | M3003 | Sigma-Aldrich | 0.3 mg | 63 | 50 ng | **0.010425** |
| **Diphosphonucleotide kinase** | N2635 | Sigma-Aldrich | 0.1 mg | 233 | 4.1 ng | **0.009517** |
| **T7 RNAP** | M0251S | New England Biolabs | 100 uL | 96 | 40 nL | **0.038360** |
| **RNAse inhibitor** | M0314S | New England Biolabs | 75 uL | 101 | 10 nL | **0.013413** |
| Total Price per 1µL |  |  |  |  |  | **0.072** |

Ribosomes
| Compound | Catalog Number | Company | Amount | Price [USD] | Amount per 1uL reaction | Price per 1uL reaction [USD] |
| --- | --- | --- | --- | --- | --- | --- |
| **Ribosomes** | M0314S | New England Biolabs | 1 mg | 250 | 3 µg | **0.815163** |
| Total Price per 1µL |  |  |  |  |  | **0.815** |

**Table S8:**
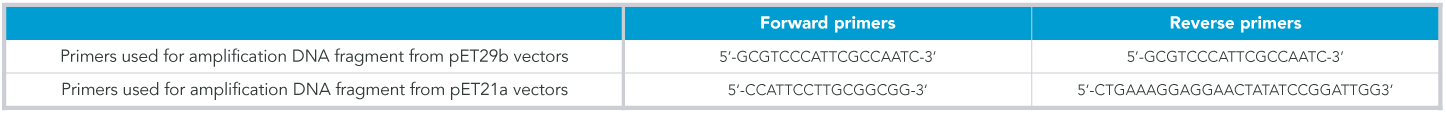
DNA sequences of primers used for CPEC

**Table S9:**
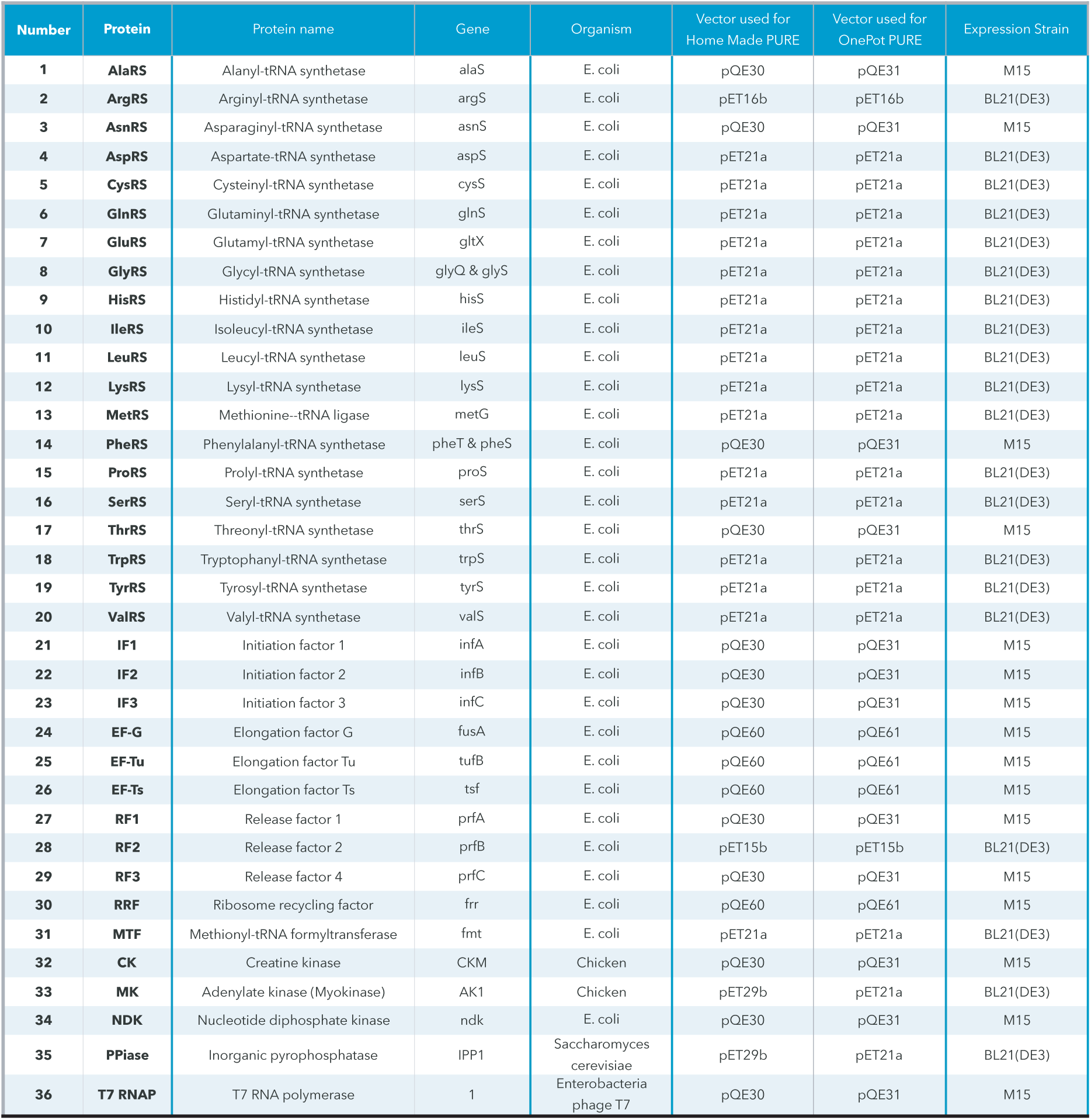
PURE protein list

**Table S10:**
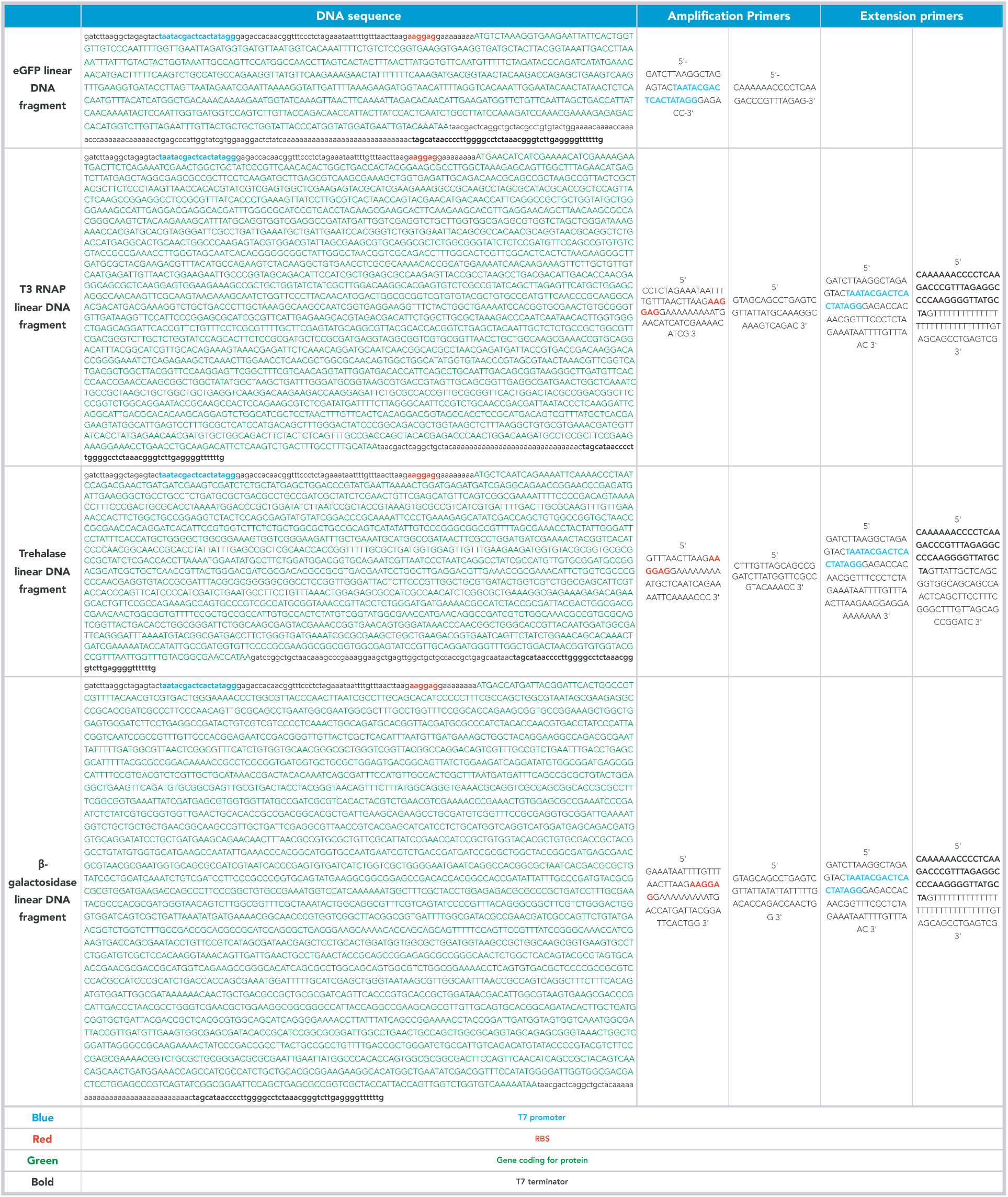
DNA sequences for linear templates with T7 RNAP promoter

**Table S11:**
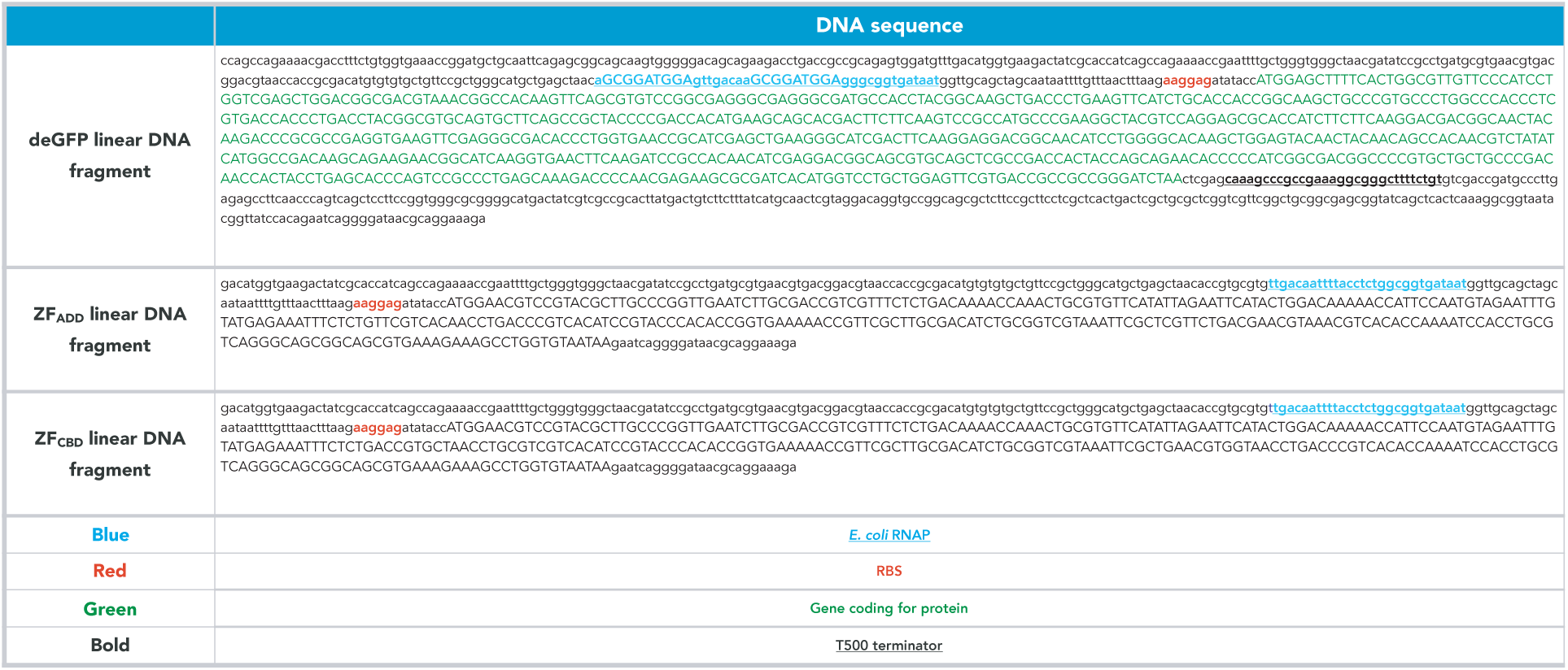
DNA sequences for linear templates with *E. coli* RNAP promoter

**Table S12:** Buffers and energy solution

Bufferes for protein purification
| Compound | Catalog number | Company | Buffer A | Buffer B | Buffer HT | Stock buffer | Stock buffer B | Note |
| --- | --- | --- | --- | --- | --- | --- | --- | --- |
|  |  |  | mM | mM | mM | mM | mM |  |
| HEPES | H0887-100ML | Sigma-Aldrich | 50 | 50 | 50 | 50 | 50 | pH = 7.6, KOH |
| Ammonium chloride | 09718-250G | Sigma-Aldrich | 1000 |  |  |  |  |  |
| Magnesium chloride | 63020-1L | Honeywell Fluka | 10 | 10 | 10 | 10 | 10 |  |
| Potassium chloride | P5405-1KG | Sigma-Aldrich |  | 100 | 100 | 100 | 100 |  |
| Imidasol | I2399 | Sigma-Aldrich |  | 500 |  |  |  | pH = 7.6, KOH |
| Glycerol | G7757-1L | Sigma-Aldrich |  |  |  | 30% | 60% |  |
| β-mercaptoethanol | M6250-100ML | Sigma-Aldrich | 7 | 7 | 7 | 7 | 7 |  |

Bufferes for ribosomes purification
| Compound | Catalog number | Company | Suspension buffer | Suspension buffer high salt | Buffer C | Buffer D | Cusion buffer | Ribosome buffer | Note |
| --- | --- | --- | --- | --- | --- | --- | --- | --- | --- |
|  |  |  | mM | mM | mM | mM | mM | mM |  |
| HEPES | H0887-100ML | Sigma-Aldrich | 10 | 10 | 20 | 20 | 20 | 20 | pH = 7.6, KOH |
| Magnesium acetate | M0631 | Sigma-Aldrich | 10 | 10 | 10 | 10 | 10 | 6 |  |
| Potassium chloride | P5405-1KG | Sigma-Aldrich | 50 | 50 |  |  |  | 30 |  |
| Ammonium chloride | 09718-250G | Sigma-Aldrich |  |  |  |  | 30 |  |  |
| Ammonium sulfate | A4418 | Sigma-Aldrich |  | 3000 | 1500 |  |  |  | pH = 7.6, KOH |
| Sucrose | 84097 | Sigma-Aldrich |  |  |  |  | 30% |  |  |
| β-mercaptoethanol | M6250-100ML | Sigma-Aldrich | 7 | 7 | 7 | 7 | 7 | 7 |  |

Energy Solution
| Compound | Catalog number | Company | Concentration in reaction | Concentration in subset (2.5x) | Units |
| --- | --- | --- | --- | --- | --- |
| Amino acids | LAA21-1KT | Sigma-Aldrich | 0.3 | 0.75 | mM |
| Magnesium acetate | M0631 | Sigma-Aldrich | 11.8 | 29.5 | mM |
| Potassium glutamate | 49601 | Sigma-Aldrich | 100 | 250 | mM |
| DTT | sc-29089B | SantaCruz Biotech | 1 | 2.5 | mM |
| ATP | R0481 | ThermoFisher | 2 | 5 | mM |
| GTP | R0481 | ThermoFisher | 2 | 5 | mM |
| CTP | R0481 | ThermoFisher | 1 | 2.5 | mM |
| UTP | R0481 | ThermoFisher | 1 | 2.5 | mM |
| tRNA | 10109541001 | Roche | 52 | 130 | U<sub>A260</sub>/mL |
| Creatine phosphate | 27920 | Sigma-Aldrich | 20 | 50 | mM |
| Folinic acid | PHR1541 | Sigma-Aldrich | 0.02 | 0.05 | mM |
| Spermidine | S2626 | Sigma-Aldrich | 2 | 5 | mM |
| HEPES | H0887-100ML | Sigma-Aldrich | 50 | 125 | mM |

## Graphical TOC Entry

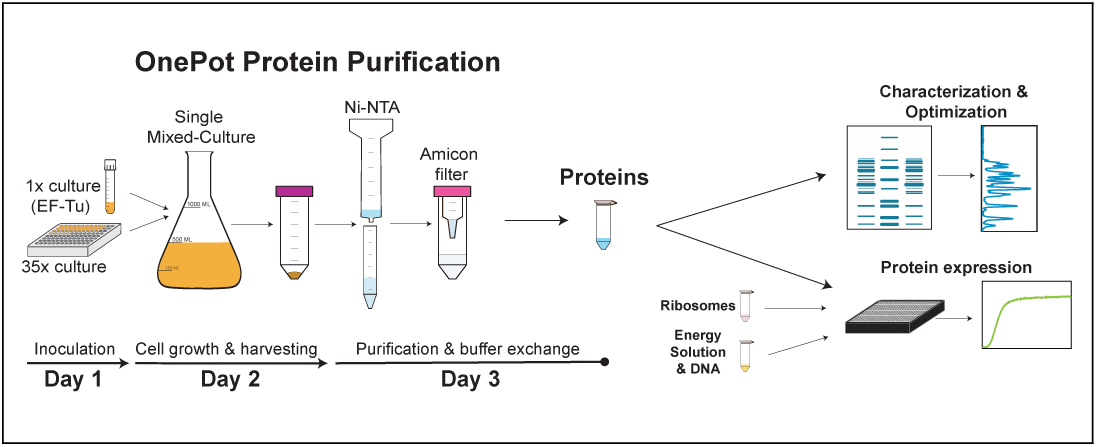

## Abbreviations

PURE: protein synthesis using recombinant elements;
Ni-NTA: nickel nitrilotriacetic acid

